# CD8^+^ T-cells protect against experimental atherosclerosis through antigen-specific interactions in the plaque

**DOI:** 10.64898/2026.11.09.765252

**Authors:** E.A. de Vlieger, M.J.M. de Jong, J. Molenaar, P.P. Lindenbergh, R.M.H. Hamelynck, K. Liu, A.C. Foks, I. Bot, B. van de Sluis, B. Slütter

## Abstract

**Background:** Atherosclerosis is a chronic inflammatory disease with a significant adaptive immune component. CD8^+^ T-cells accumulate in plaques and display signatures of antigen-specific activation. However, whether this accumulation is driven by local antigen recognition and what functional consequences follow, remains poorly understood. Here we address this gap of knowledge using loss- and gain-of-function experiments.

**Methods:** Atherosclerotic *Rag^-/-^Ldlr*^-/-^ mice received adoptively transferred wildtype or antigen-restricted CD8^+^ T-cells alongside CD4^+^ T-cells to distinguish antigen-specific from antigen-independent responses. In a complementary model, *Apob^OVA-TG^xLdlr^-/-^*mice – expressing an ovalbumin-tagged variant of the atherosclerosis-associated ApoB_100_ protein – enabled OT-I CD8^⁺^ T-cells to function as atherosclerosis-antigen-specific CD8^⁺^ T-cells. T-cell distribution, activation and phenotype were assessed systemically and within lesions, plaque burden and composition were quantified histologically.

**Results:** Wildtype CD8^+^ T-cells accumulated significantly more in aortic lesions compared to antigen-restricted CD8^+^ T-cells, and exhibited increased activation, exhaustion and cytokine secretion, indicating that plaque-infiltrating T-cells are predominantly antigen-specific. Perturbing antigen-specific CD8^+^ T-cell interactions significantly decreased plaque stability index, suggesting a protective role for antigen-specific CD8^+^ T-cells. In *Apob^OVA-TG^xLdlr^-/-^*mice, OT-I T-cells accumulated at high frequency in the liver, consistent with hepatic ApoB_100_-expression, and were found in circulation and aorta. Histological analysis revealed reduced plaque burden in OT-I recipients, reproducing an atheroprotective effect of antigen-specific CD8^+^ T-cell responses. Plaque reduction coincided with reduced TUNEL, but similar Caspase 3 staining, suggesting OT-I transfer results in reduced secondary necrosis. In line with this, CD47 expression in OT-I recipient mice was reduced and *in vitro*, OT-I, but not antigen non-specific CD8^+^ T-cells reduced expression of the “don’t eat me”-signal CD47 on OVA-presenting foam cells.

**Conclusions:** CD8^+^ T-cell accumulation in experimental atherosclerosis is driven primarily by antigen-specific interactions and such responses are associated with plaque stabilization and reduced lesion formation, potentially through induction of apoptosis and prevention of secondary necrosis.

## Introduction

Cardiovascular disease remains the leading cause of morbidity and mortality worldwide, with atherosclerosis as its primary underlying pathology.^1^ Atherosclerosis is a chronic, lipid-driven inflammatory disease characterized by the subendothelial accumulation of oxidized low-density lipoprotein (oxLDL), foam cell formation and progressive plaque development. Although lipid-lowering therapies reduce cardiovascular risk, a persistent inflammatory burden in treated patients underscores the need to better understand the immune mechanisms driving plaque progression and destabilization.^2-4^

Next to innate immune cells, adaptive immune responses have emerged as important modulators of atherogenesis, with T-cells playing a central role.^5, 6^ Among these, CD8^+^ cytotoxic T-cells have gained increasing attention for their capacity to drive tissue inflammation through direct cytotoxicity, IFN-γ and TNF-α production, and modulation of macrophage and smooth muscle cell behavior.^7^ CD8^+^ T-cells are present in both human and murine atherosclerotic plaques,^8-11^ and studies in mouse models have reported both pro- and anti-atherogenic effects depending on experimental context,^7, 12-15^ reflecting the complexity of their functional roles in plaque development.

The contrasting results of CD8^+^ T-cells in experimental atherosclerosis may be related to their diverse T-cell receptor (TCR) repertoire. Recent single-cell analysis has demonstrated the presence of clonally expanded CD8^+^ T-cells in human atherosclerotic lesions,^9, 11^ with a substantial CD8^+^ T-cell population also identified in murine plaques.^8^ Of note, clonal expansion was less pronounced in plaque-derived CD8^+^ T-cells compared to their circulating counterparts, raising the question of whether these T-cells accumulate locally in response to plaque-specific antigens, or are instead recruited through antigen-independent inflammatory signaling.^11^ Supporting local activation, plaque-derived CD8^+^ T-cells exhibited elevated CD69 expression, a marker of tissue-resident and recently activated T-cells.^11, 16^ However, whether this activated phenotype reflects true antigen-driven stimulation or bystander activation remains unknown. Indeed, our recent work demonstrated that virus-specific CD8^+^ T-cells in atherosclerotic plaques display an equally activated phenotype as other plaque-derived CD8^+^ T-cells, despite the absence of cognate viral antigen in the plaque microenvironment, suggesting that antigen-independent mechanisms may substantially contribute to CD8^+^ T-cell activation *in situ.*^17^

Resolving whether CD8^+^ T-cell accumulation in plaques is antigen-driven and whether modifying atherosclerosis-specific CD8^+^ T-cell responses affects disease progression has direct therapeutic relevance. Here, we address these questions using two complementary murine models, one in which antigen-specific CD8^+^ T-cell interactions are perturbed and one in which they are deliberately induced. Supported by macrophage-CD8^+^ T-cell co-culture experiments, these models allow us to dissect antigen-specific and antigen-independent contributions of CD8^+^ T-cells to plaque development and evaluate their potential as targets for therapeutic intervention.

## Methods

### Mice

C57BL/6J, C57BL/6-Tg(TcraTcrb)1100Mjb/J (OT-I) mice, and *Ldlr*^−/−^ mice were purchased from Jackson Laboratory (Bar Harbor, Maine, USA) and bred in-house. *Rag1^-/-^Ldlr^-/-^* mice and *Rag1^-/-^OT-I^+/+^*mice were crossed and bred in-house. Mice were housed under standard laboratory conditions, with a 12-hour dark/light cycle and food and water were provided ad libitum. All animal work was performed in compliance with the Dutch government guidelines and the Directive 2010/63/EU of the European Parliament and experiments were approved by the Ethics Committee for Animal Experiments and the Animal Welfare Body of Leiden University. Prior to each experiment, mice were randomized based on age, weight and cholesterol levels. Experiments were performed in a blinded fashion, only EV and MJ were aware of assigned treatments.

### Generation of Apob^OVA-TG^ mice

*Apob*^OVA-TG^ mice were generated by inserting the DNA sequence encoding the immunodominant peptides of ovalbumin (OVA_257_ and OVA_323_) before the stop codon of the *Apob* gene using CRISPR/Cas9-mediated gene editing technology as previously described.^18^ C57BL/6J were superovulated by injection with 5 IU Folligonan (0.2 mL i.p.), and 46 hours later with 5 IU Chorulon (0.2 mL i.p.), whereafter they were housed with fertile C57BL/6J mice. The next day, zygotes were isolated from the oviduct and micro-injected with 100 ng/ul *Cas9* RNA, 50 ng/µL sg*Apob* (TCTGATTCATCTACTAACAA) and 100 ng/µL repair oligo. Injected zygotes were transferred to the oviduct of pseudopregnant females at the same day. Offspring carrying the correct modification, identified by PCR with two primers flanking the insertion site (Primer 1: TTGGTGAGTCCACAAGATTGA, Primer 2: TGTGCTTCTTGCTCTTGCTG, wildtype = 268bp, *Apob^OVA-TG^* = 368bp), were selected for breeding with *Ldlr*^-/-^ mice. Germline transmission of the mutation was verified and the mice carrying the modification were backcrossed into the *Ldlr*^-/-^ mice, resulting in *Apob^OVA-TG^* mice exhibiting a heterozygous *Apob^OVA-TG^* modification and a homozygous *Ldlr* knockout.

### Western blotting

To validate the expression of the OVA-tag at the protein level, a Western blot was performed. In brief, 10 µL of denatured serum was loaded into a 4–20% precast polyacrylamide gel (Bio-Rad), and electrophorized for 1.5 hours at 120V, followed by 1.5 hours of blotting at 200mA. Blots were primarily stained with a biotinylated anti-OVA_323_ antibody (1:1000, Innovagen), specifically binding one of the inserted OVA peptides, and secondarily stained with streptavidin-HRP (1:5000, BD Biosciences). HRP was activated by adding chemiluminescent substrate (SuperSignal West Pico Chemiluminescent Substrate) and signal was imaged using an Amersham Imager 600. To confirm the OVA-tag was inserted to ApoB_100_, blots were stripped of the OVA_323_ antibody and stained with a biotinylated anti-ApoB_100_ antibody (1:500, Bio-Techne), followed by secondary staining and imaging as described above.

### Adoptive transfer of naïve antigen-specific T-cells into Apob^OVA-TG^ mice

5 female *Apob^OVA-TG^* mice or 5 *Ldlr^-/-^* mice between 8-15 weeks of age received 1·10^6^ ovalbumin specific OT-II CD4^+^ T-cells and OT-I CD8^+^ T-cells. OT-II CD4^+^ and OT-I CD8^+^ T-cells were isolated from the spleens of CD45.1-expressing OT-I/OT-II mice by negative selection using CD4^+^ or CD8^+^ T-cell isolation kits (Miltenyi Biotec), according to the manufacturer’s instructions. Upon transfer, mice were fed a Western-type diet for 6 weeks to induce lesion formation. OT-I/OT-II donor cells were identified based on their expression of CD45.1. Mice with an OT-I/OT-II population of >0.03% of the total CD4^+^/CD8^+^ T-cell population were included in the phenotypic analysis.

### Adoptive transfer of antigen-specific or antigen-independent T-cells

29 female *Rag1^-/-^Ldlr*^-/-^ mice between 8-15 weeks of age received a combination of either 6·10^5^ wildtype CD4^+^ and 3·10^5^ wildtype CD8^+^ T-cells or 3·10^5^ *Rag1^-/-^Ldlr^-/-^OT-I^+/+^*CD8^+^ T-cells. Transferred CD4^+^ and CD8^+^ T-cells were isolated from spleens using negative selection CD4^+^ or CD8^+^ T-cell isolation kits (MojoSort™ Mouse CD4^+^ or CD8^+^ T-cell, BioLegend). Upon transfer, mice were fed a Western-type diet for 10 weeks to induce lesion formation.

### Adoptive transfer of activated antigen-specific CD8^+^ T-cells

30 female *Apob^OVA-TG^xLdlr^-/-^*mice between 8-20 weeks of age received either *ex vivo* activated 6·10^6^ OT-I CD8^+^ T-cells or PBS control. Transferred CD8^+^ T-cells were isolated from spleens using negative selection CD8^+^ T-cell isolation kits (MojoSort™ Mouse CD4^+^ or CD8^+^ T-cell, BioLegend). Following negative selection, CD8^+^ T-cells were cultured overnight (500,000 cells/mL) in complete RPMI (cRPMI, RPMI medium (VWR) supplemented with 10% fetal bovine serum (FBS, Merck), 1% penicillin/streptomycin (Fisher Scientific), 1% L-glutamine (VWR), and 0.05 mM β-mercaptoethanol) containing anti-CD3 (1 µg/mL, eBioscience), anti-CD28 (1 µg/mL, eBioscience) and IL-2 (200 U/mL, BioLegend) at 37°C and 5% CO_2_. The following day, the culture medium was replaced with fresh cRPMI supplemented with IL-2 (200 U/mL, BioLegend), and cultured overnight before intravenous adoptive transfer via the tail vein. Recipient mice were then fed a Western-type diet for 7 weeks. CD8^+^ T-cell populations and cholesterol levels were monitored throughout the study.

### Sacrifice

At the end of the experiments, mice were anesthetized intraperitoneally with a mix of ketamine (100 mg/mL) and Xylazine (10 mg/mL). Blood was collected via orbital bleeding, followed by *in situ* perfusion with saline through the left ventricle. Heart, aorta, heart lymph nodes, inguinal lymph nodes, liver and spleen were collected for further analysis.

### Organ processing

Serum was obtained from whole blood collected in clot-inducing Microvette CB300 Z tubes (Sarstedt) and separated by centrifugation at 8,000 rpm for 10 minutes at 4°C. Serum was stored at -80°C until analysis. Single-cell suspensions from spleens and lymph nodes were obtained by passing the tissues through 70 μm cell strainers (Greiner Bio-One). Red blood cells were lysed in spleen and blood samples using ACK lysis buffer (0.15M NH_4_Cl, 1mM KHCO_3_, 0.1mM Na_2_EDTA, pH 7.3) for 2 min at room temperature. The upper portion of the heart was embedded in Tissue-Tek OCT (Sakura) and stored at -80°C until further use. Liver was mashed through 70 μm cell strainer and subsequently digested for 30 min at 37 °C under constant agitation by incubation with collagenase I (450 U/mL) collagenase XI (250 U/mL) and DNAse (120 U/mL) (all Sigma-Aldrich). Aortas were cut into approximately 1 mm^2^-fragments and digested for 30 min at 37 °C under constant agitation in digestion buffer containing collagenase I (450 U/mL) collagenase XI (250 U/mL), DNAse (120 U/mL) and hyaluronidase (120 U/mL) (all Sigma-Aldrich). Digested tissue was then passed through a 70-μm cell strainer to obtain single-cell suspensions.

### Cholesterol assay

Serum total cholesterol levels were assessed through an enzymatic colorimetric analysis (Roch/Hitachi, Germany), using Precipath standardized serum (1.69 mg/ml, Roche/Hitachi) as an internal standard.

### Alanine aminotransferase (ALT) activity assay

Serum ALT activity was determined using the ALT Activity Assay Kit (MAK-052, Sigma-Aldrich) following the manufacturer’s protocol. ALT activity was expressed as milliunits per milliliter (mU/mL), where one unit of ALT activity is defined as the amount of enzyme that generates 1 µmol of pyruvate per minute under assay conditions.

### Flow cytometry

Flow cytometry was performed on ∼5·10^5^ cells, or all available cells, by staining with the appropriate antibodies (Supplementary Table 1), TruStain FcX (anti-mouse CD16/32) Antibody (BioLegend), and live/dead viability staining (eBioscience and BioLegend) in PBS containing 2% FCS, according to manufacturer’s instructions. For intracellular staining, cells were fixed and permeabilized using either the Intracellular Fixation and Permeabilization Buffer Set or the FOXP3/Transcription Factor Staining Buffer Set (eBioscience), as appropriate. Flow cytometry was performed on a Cytoflex S (Beckman Coulter) or Cytek Northern Lights (V/B/R, 3-laser) and analyzed using FlowJo v10.8 software (BD Life Sciences). Gating strategy for cytokine staining is shown in Figure S5.

### Histological analysis

Cryosections (7 µm) of the trivalve region of the heart were prepared using a Leica CM1950 cryostat. Plaque size and stenosis were assessed by Oil Red O staining (Sigma-Aldrich), whereas collagen content and necrotic core area were evaluated by Masson’s Trichrome staining (Sigma-Aldrich). Intralesional macrophages were detected by immunohistochemistry using anti-MOMA-2 (1:1,000, Bio-Rad) followed by biotinylated rabbit anti-rat IgG (1:200, Vector) and ImmPACT NovaRED (Vector). Apoptotic cells were detected by TUNEL staining (HY-K1091A, MedChemExpress) according to manufacturer’s instructions. Caspase-3 positive area was detected by immunohistochemistry using Cleaved Caspase 3 polyclonal antibody (1:1,600, Proteintech) followed by BrightVision goat anti-rabbit IgG HRP (Immunologic) and ImmPACT NovaRED (Vector). CD47 positive area was detected by immunohistochemistry using CD47 polyclonal antibody (1:8,000, Proteintech) followed by BrightVision goat anti-rabbit IgG HRP (Immunologic) and ImmPACT NovaRED (Vector). Sections were digitized using a Panoramic 250 Flash III (3DHISTECH), ZEISS scanner or Leica DM-RE microscope and analyzed with QuPath.^19^ Plaque area was quantified from five serial sections per mouse, all other histological analyses were performed on three consecutive sections. Hearts lacking a trivalve area or damaged hearts were excluded.

### In vitro macrophage-CD8^+^ T-cell coculture

Bone-marrow derived macrophages (BMDMs) were generated from bone marrow from the femurs and tibias of 8-10 week old male C57BL/6J mice. Bone marrow cells were cultured at 1·10^6^ cells/mL in cRPMI supplemented with M-CSF (10 ng/mL, BioLegend) for 7 days, with medium replaced on day 3. On day 7, adherent BMDMs were harvested and seeded into F-bottom 96-well plates at 5·10^4^ cells/well and exposed to oxidized LDL (20 µg/mL, Invitrogen™) overnight, followed by overnight pulsing with OVA257 peptide (1 µM, Invivogen). CD8^+^ T-cells were isolated from the spleens of C57BL/6J or OT-I mice by negative selection using the MojoSort Mouse CD8^+^ T-cell Isolation Kit (BioLegend). CD8^+^ T-cells were cocultured with BMDMs at a macrophage cell ratio of 1:2 (5·10^4^ BMDMs and 1·10^5^ CD8). Macrophage marker expression was assessed by flow cytometry after 48 hours of co-culture.

### Single-cell RNA sequencing human atherosclerotic carotid plaques

Publicly available human plaque datasets (GSE155512^20^, GSE159677^21^, GSE253903^22^) were downloaded from Gene Expression Omnibus (GEO) and combined into a single Seurat object in R (version 4.4.1) using Seurat package (version 5.1.0^23^). Quality control was carried out by excluding cells with a percentage of mitochondrial gene expression of less than 10%, a count of fewer than 500 unique genes or a total count of less than 800. Double cells and ambient RNA contamination were removed using scDblFinder (version 1.18.0^24^) and the decontX function of the celda R package respectively (version 1.20.0^25^), whereafter a total of 100,658 cells remained, of which 19,955 contained CITE-seq data. The object was split into patient-specific layers, whereafter normalization, scaling and variable features identification was performed on each layer individually using SCTransform. Subsequently, the IntegrateLayers function was applied for integration, specifying ‘HarmonyIntegration’ as the ‘method’ argument and ‘SCT’ as the ‘normalization.method’ argument. ScICE was used for the cell clustering.^26^ Seurat Harmony reductions and UMAP coordinates were used as input for the Julia-based scLENS pipeline^27^ to calculate cluster inconsistency coefficients and obtain consistent cluster labels. Plots were generated using the ggplot2 package (3.5.20^28^).

### Athero-Express Biobank gene comparison

Bulk RNA-sequencing data were analyzed from 1093 carotid atherosclerotic plaques from the Athero-Express Biobank.^29^ The expression levels of *CD47, CD3E, CD4, CD8A* and *CD8B* were compared using the Gene Lookup & Expression Explorer available at www.atheroportal.nl. Pairwise Spearman correlations were performed.

### Statistical analysis

Animal group sizes were determined based on group means, with vascular occlusion as the primary outcome measure. Data were analyzed using GraphPad Prism (version 10, GraphPad Software). Outliers were identified using the ROUT method and normality was assessed with the Shapiro-Wilk test. Normally distributed data were compared using Student’s t-test or one-way ANOVA with Tukey’s multiple-comparisons test, depending on group number, non-normally distributed data were compared using the Mann-Whitney U-test or Kruskal-Wallis test. All data is presented as mean±SD.

## Results

### Antigen-driven CD8^+^ T-cell accumulation in atherosclerotic plaques

CD8^+^ T-cells are generally recruited towards sites of inflammation in an antigen-independent fashion, based on recognition of activated endothelium and chemokine receptor signaling, including CXCR4.^30^ Yet retention of CD8^+^ T-cells in inflamed tissue may rely on antigen recognition, with TCR-driven expression of CD69 being a major cue for tissue residence. To address whether antigen recognition plays a role in the accumulation of CD8^+^ T-cells in the atherosclerotic plaque, we developed an experimental model that excludes antigen-driven accumulation of CD8^+^ T-cells in atherosclerotic plaques. *Rag1*-deficient (*Rag1^-/-^*), which lack T- and B-cells, were crossed with *Ldlr*-deficient (*Ldlr^-/-^*) mice and received a Western-type diet to induce atherosclerosis. All mice received an adoptive transfer of CD4^+^ T-cells from wildtype (WT) C57BL/6J mice and an adoptive transfer of CD8^+^ T-cells from WT C57BL/6J or *Rag1^-/-^ xOT-I* TCR transgenic mice, and Western-type diet was continued for another 10 weeks (Figure 1A).

**Figure 1.**
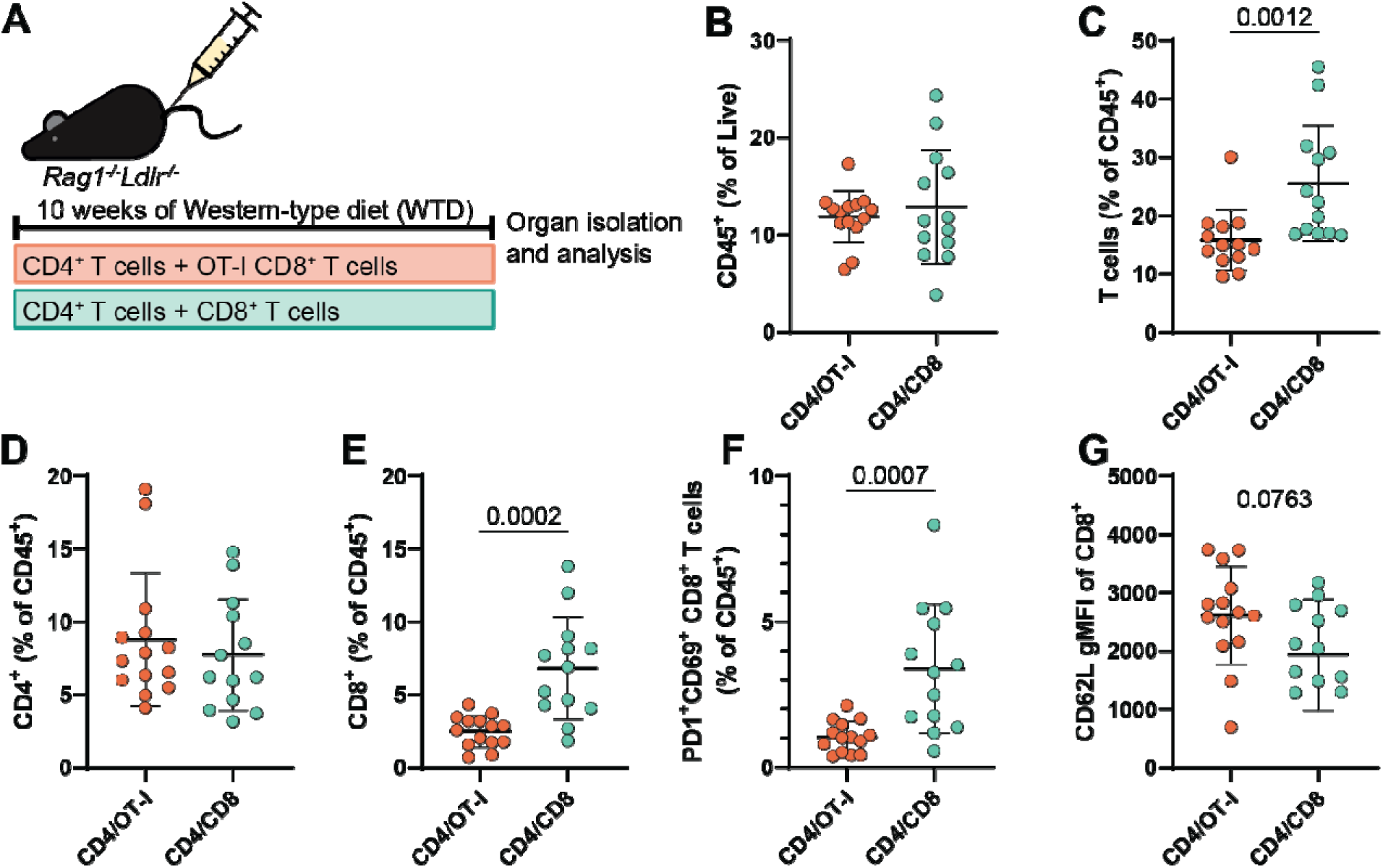
CD8^+^ T-cells accumulate in the aorta in antigen-specific manner upon development of atherosclerosis. (A) Experimental setup of adoptive transfer of WT or OT-I T-cells into Rag1^-/-^Ldlr^-/-^ mice. The proportion of (B) CD45^+^ immune cells of live cells, and (C) Thy1.2^+^ T-cells, (D) CD4^+^ T-cells, (E) CD8^+^ T-cells, and (F) PD1^+^CD69^+^CD8^+^ T-cells of CD45^+^ immune cells in the aortic arch after 10 weeks of Western-type diet. (G) Geometric mean CD62L expression in CD8^+^ T-cells. Outliers were removed using the ROUT test, followed by Shapiro-Wilk normality testing. Based on normality, unpaired t-tests or Mann-Whitney tests were performed. n=13-14 mice per group. Data are shown as mean±SD.

Total CD45^+^ aortic immune cell numbers were comparable between both groups (Figure 1B), while the proportion of T-cells was significantly increased in WT T-cell recipients compared to antigen-restricted OT-I recipients (Figure 1C). This effect is due to a 63% reduction in CD8^+^ T-cell accumulation in the antigen-restricted group, as CD4 T-cell levels did not differ between groups (Figure 1D-E). Importantly, reduced accumulation of OT-I T-cells in the aorta was not explained by a general reduction of these T-cells in the mice, as systemically similar numbers of CD8^+^ T-cells were detected in both groups (Figure S1A,F,K,P). Within the aorta of WT T-cell recipients, an accumulation of PD1^+^CD69^+^ CD8^+^ T-cells was observed (Figure 1F), a phenotype associated with local antigen-driven activation.^31^ Additionally, WT CD8^+^ T-cells showed a trend toward reduced CD62L expression compared to OT-I CD8^+^ T-cells, all suggestive of CD8^+^ T-cells undergoing antigen-driven retention within the plaque. Systemically, WT CD8^+^ T-cells displayed a more effector-memory phenotype in the blood, spleen, inguinal lymph node (iLN) and mediastinal lymph node (mLN) than their antigen-restricted counterparts (Figure S1). In the mLN which drains the peritoneal cavity and its vasculature including the aorta, WT CD8^+^ T-cells cells also showed increased PD1 and Granzyme B expression compared to the antigen-restricted CD8^+^ T-cell transfer group (Figure S1S,T). Overall, this experiment strongly suggests that CD8^+^ T-cells in murine atherosclerotic plaques are dependent on cognate antigens to accumulate in the lesion and can undergo local activation.

Having established that CD8^+^ T-cells accumulate in aortic lesions in an antigen-driven manner, we next investigated whether this was accompanied by functional differences in effector capacity. CD8^+^ T-cells isolated from the mLN and spleen were stimulated *ex vivo* with PMA/ionomycin for 6 hours and assessed for cytokine and granzyme production. WT CD8^+^ T-cells exhibited significantly greater capacity to produce IFN-γ (Figure 2A,F) and Granzyme B (Figure 2B,G) compared to OT-I T-cells in both organs, indicating that antigen-specific CD8^+^ T-cells can adopt a more pronounced cytotoxic effector phenotype. Notably, WT CD8^+^ T-cells in the mLN were also significantly more capable of producing IL-4 than OT-I T-cells, an effect that was not observed in the spleen (Figure 2C,H). Although IL-4 is classically associated with Th2 CD4 T-cell responses, its production by CD8^+^ T-cells has been reported in the context of chronic antigen stimulation^32, 33^ and its slight elevation in the mLN may reflect a local response to antigen exposure in the lymph node close to the heart and aorta. WT CD8^+^ T-cells in the mLN also secreted less IL-10 compared to OT-I T-cells (Figure 2D), with no difference observed in the spleen (Figure 2I), suggesting that the reduction in this anti-inflammatory cytokine is selective to the areas close to the atherosclerotic lesions. Finally, IL-17, a cytokine linked to neutrophil recruitment and vascular inflammation,^34^ was significantly elevated in WT CD8^+^ T-cells from the spleen, though at a low percentage (Figure 2J). A similar though non-significant trend occurs in the mLN (Figure 2E), suggesting a role for antigen-specific CD8^+^ T-cells in increasing local inflammatory signaling beyond the lesion. Importantly, cytokine levels were similar between groups for the CD4 T-cell compartment (Figure S2), showing the specificity of our experimental set-up.

**Figure 2.**
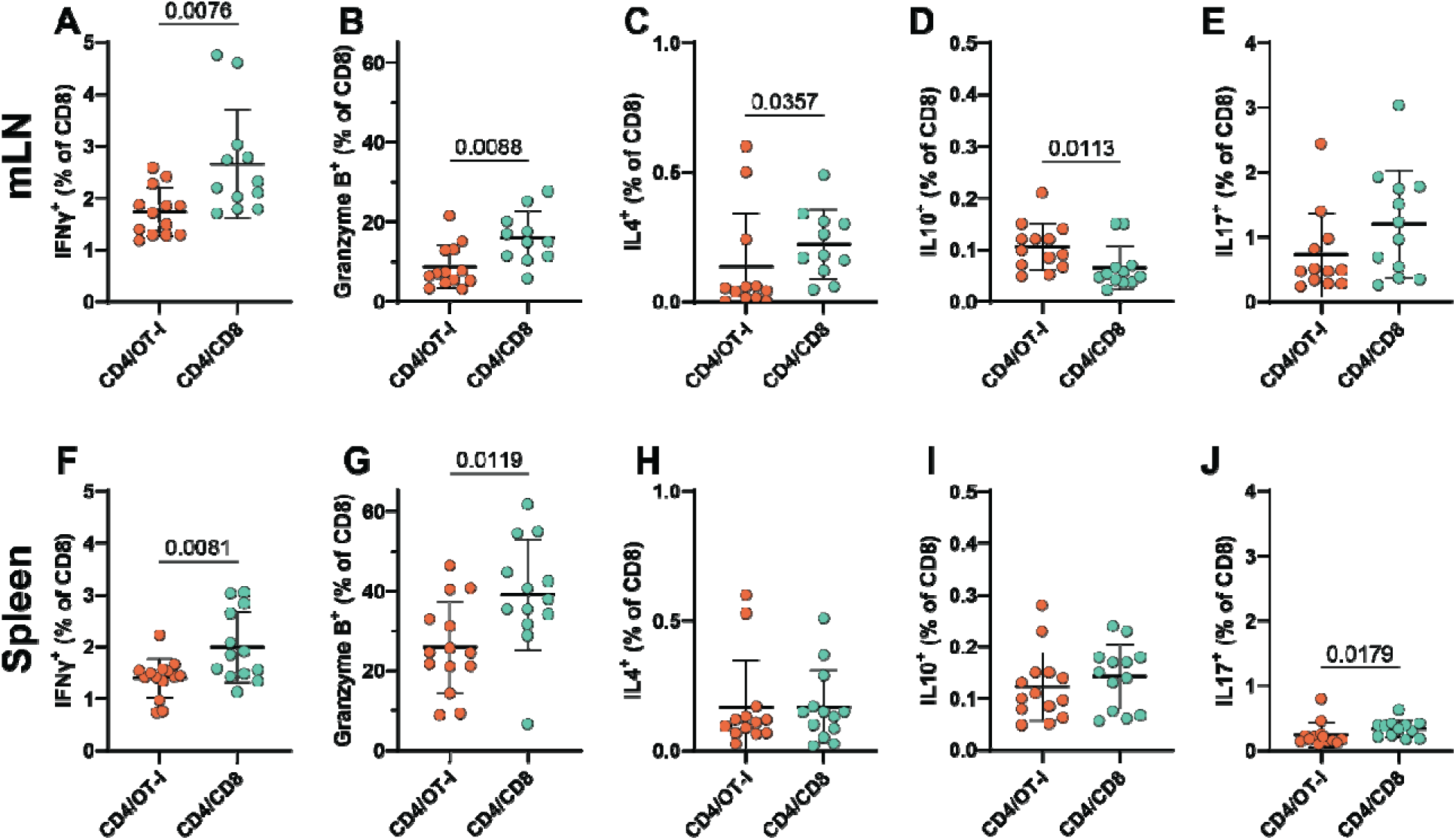
Antigen-specific CD8^+^ T-cells are more activated than antigen-restricted CD8^+^ T-cells. Percentage of CD8^+^ T-cells producing cytokines and granzymes in heart lymph nodes (A-E) and spleen (F-J) after 6 hours of ex vivo PMA/ionomycin stimulation. Interferon gamma (A,F), Granzyme B (B,G), interleukin-4 (C,H), interleukin-10 (D,I) and interleukin-17 production (E,J) in CD8^+^ T-cells. Outliers were removed using the ROUT test, followed by Shapiro-Wilk normality testing. Based on normality, unpaired t-tests or Mann-Whitney tests were performed. n=13-14 mice per group. Data are shown as mean±SD.

Together, enhanced inflammatory cytokine and Granzyme B production by WT CD8^+^ T-cells compared to antigen-restricted CD8^+^ T-cells is most prominent in the proximity of the atherosclerotic plaques (mLN), but its activation is not limited to the plaque and is also reflected by systemic CD8^+^ T-cell responses.

### Absence of antigen-specific CD8^+^ T-cells destabilizes murine atherosclerotic lesions

Next, we address whether an absence of antigen-specific CD8^+^ T-cells translates into differences in plaque composition and stability (Figure 3). Lesion size and composition were assessed in the aortic root. The vascular occlusion in the three-valve area was comparable between WT and OT-I recipients (Figure 3A), indicating that antigen-specific CD8^+^ T-cell responses do not significantly affect overall lesion burden at this stage. However, qualitative differences in plaque compositions were seen. Surprisingly, WT CD8^+^ T-cell recipients showed a slight decrease in necrotic core and slight increase in collagen content, collectively resulting in a significantly higher plaque stability index compared to OT-I recipients (Figure 3B-D). Since macrophages are known to contribute to necrotic core content and unstable plaques, a reduction in myeloid infiltration could account for the observed improved plaque stability. In support of this, WT recipients displayed reduced myeloid cell percentages and absolute counts in the aortic arch compared to OT-I recipients (Figure 3G-H). Remarkably, MOMA-2 staining revealed no difference in macrophage content within the three-valve area between groups (Figure 3F), suggesting that the reduction in myeloid infiltration is either spatially restricted to the aortic arch, or that changes in macrophage viability instead of absolute numbers underlie the observed morphological differences. In support of the latter, TUNEL staining demonstrated a significant increase in apoptotic and necrotic cells in antigen-restricted OT-I recipients compared to WT recipients (Figure 3I).

**Figure 3.**
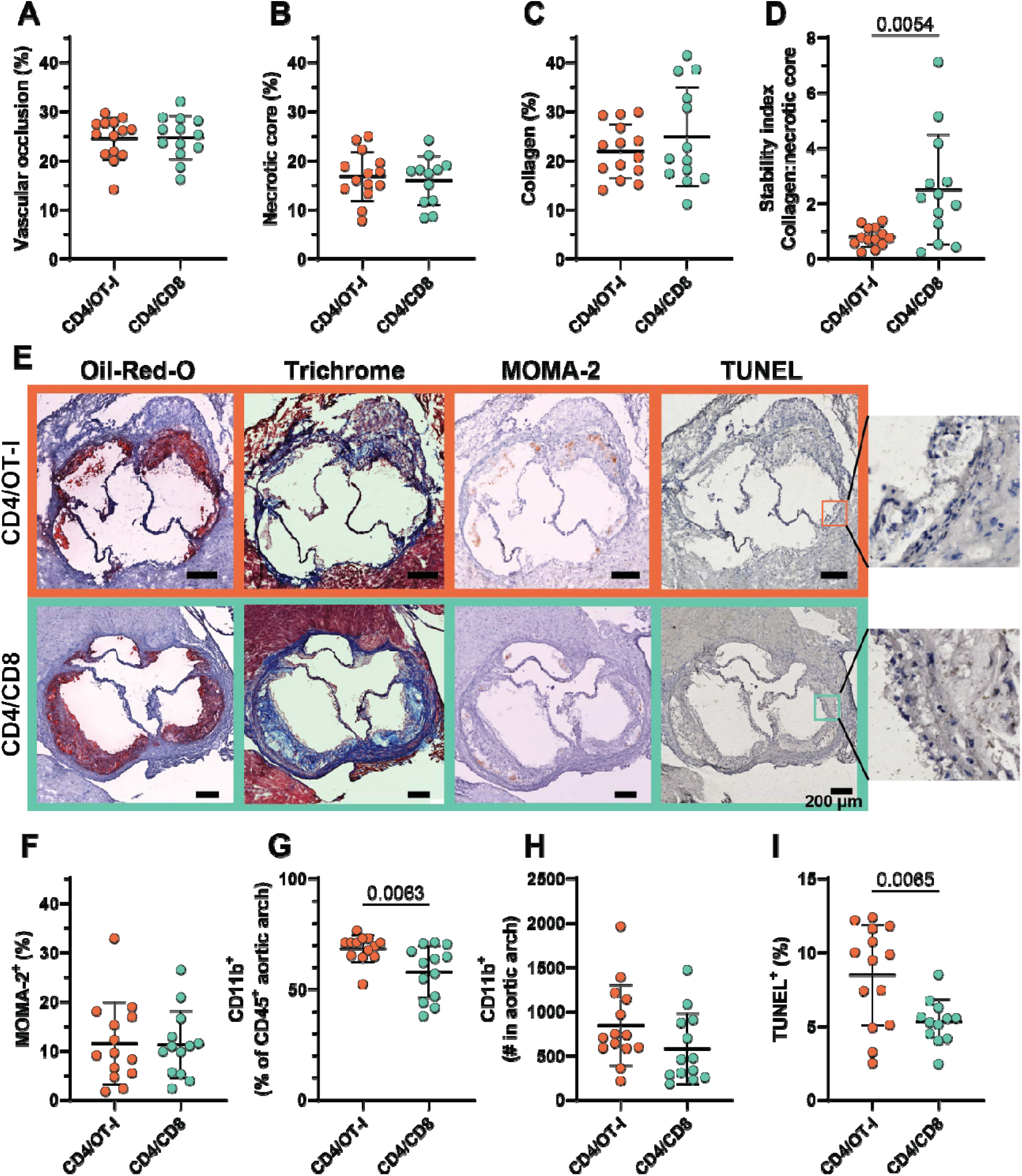
Antigen-specific CD8^+^ T-cell transfer induces more stable plaques by reducing necrotic core formation, inducing collagen retainment and reducing apoptosis. (A) Vascular occlusion in the aortic roots by Oil-red O staining. (B) Necrotic core and (C) collagen content by Masson’s Trichrome staining. (D) Stability index of plaques, by ratio of collagen per necrotic core. (E) Representative pictures of stained lesions. (F) Monocyte/macrophage content determined by MOMA-2 staining. The proportion of (G) CD11b^+^ myeloid cells of live CD45^+^ immune cells and the count of (H) CD11b^+^ myeloid cells in the aortic arch. (I) TUNEL^+^ area in *lesions as determined by TUNEL staining. Outliers were removed using the ROUT test, followed by Shapiro-Wilk normality testing. Based on normality, unpaired t-tests or Mann-Whitney tests were performed. n=13-14 mice per group. Data are shown as mean±SD*.

### ApoB-reactive CD8^+^ T-cells reduce atherosclerosis

As the absence of antigen-specific CD8^+^ T-cells disrupts plaque stability, we next sought to investigate whether enhancing their numbers could promote lesion stabilization. Apolipoprotein B 100 (ApoB_100_) is the best-characterized T-cell antigen in atherosclerosis.^35, 36^ Roy et al. recently showed that stimulation of human PBMCs with ApoB_100_ peptides elicits IFNγ-producing CD8^+^ T-cell responses, mediated through defined HLA-restricted epitopes.^37^ While this confirms the relevance of ApoB_100_-specific CD8^+^ T-cell responses in humans, the murine MHC-I-restricted epitopes that would be recognized *in vivo* remain unknown. To enable controlled introduction of atherosclerosis-antigen-specific CD8^+^ T-cell responses in mice, we developed a novel mouse model in which ApoB_100_ is genetically fused to two immunodominant domains of ovalbumin (OVA) to allow recognition of ApoB_100_-OVA by OVA-specific OT-I CD8^+^ T-cells, thereby inducing antigen-specific interactions in sites where ApoB_100_ is presented. We crossed these mice on an *Ldlr^-/-^* background and fed a Western-type diet for 7 weeks to characterize these mice. *Apob^OVA-TG^xLdlr*^-/-^ and *Ldlr*^-/-^ showed similar weight, serum cholesterol levels and atherosclerotic plaque size and stability (Figure S3). Moreover, we confirmed that serum from *Apob^OVA-TG^xLdlr*⁻*^/^*⁻^-^ mice and not serum from *Ldlr*^-/-^ mice can induce antigen-specific CD8^+^ T-cell activation in *ex vivo* cultured OT-I splenocytes (Figure S4D).

To assess the effect of antigen recognition by CD8^+^ T-cells, we transferred a limited number (1·10^6^) activated OT-I CD8^+^ T-cells into *Ldlr*^-/-^ or *Apob^OVA-TG^xLdlr^-/-^*mice and expansion was monitored during Western-type diet (Figure 4A). After six weeks a larger number of OT-I cells was detected in the mLN of *Apob^OVA-TG^xLdlr*^-/-^ mice compared to *Ldlr*^-/-^ mice lacking the cognate antigen (Figure 4B), with a similar trend observed in the spleen (Figure 4E), suggesting that OT-I T-cells expanded in the mice in response to their cognate antigen. Consistent with an antigen-specific response, splenic OT-I T-cells from *Apob^OVA-TG^xLdlr*^-/-^ mice showed increased CD69 expression, though this was not observed in the mLN (Figure 4C,F). Phenotypic characterization revealed that OT-I cells in *Apob^OVA-TG^xLdlr*^-/-^ mice adopted an effector-memory phenotype, while those in *Ldlr*^-/-^ mice remained naïve or central memory (Figure 4D,G). This indicates that ApoB-derived OVA peptides are presented *in vivo* and drive activation and effector differentiation of antigen-specific CD8^+^ T-cells.

**Figure 4.**
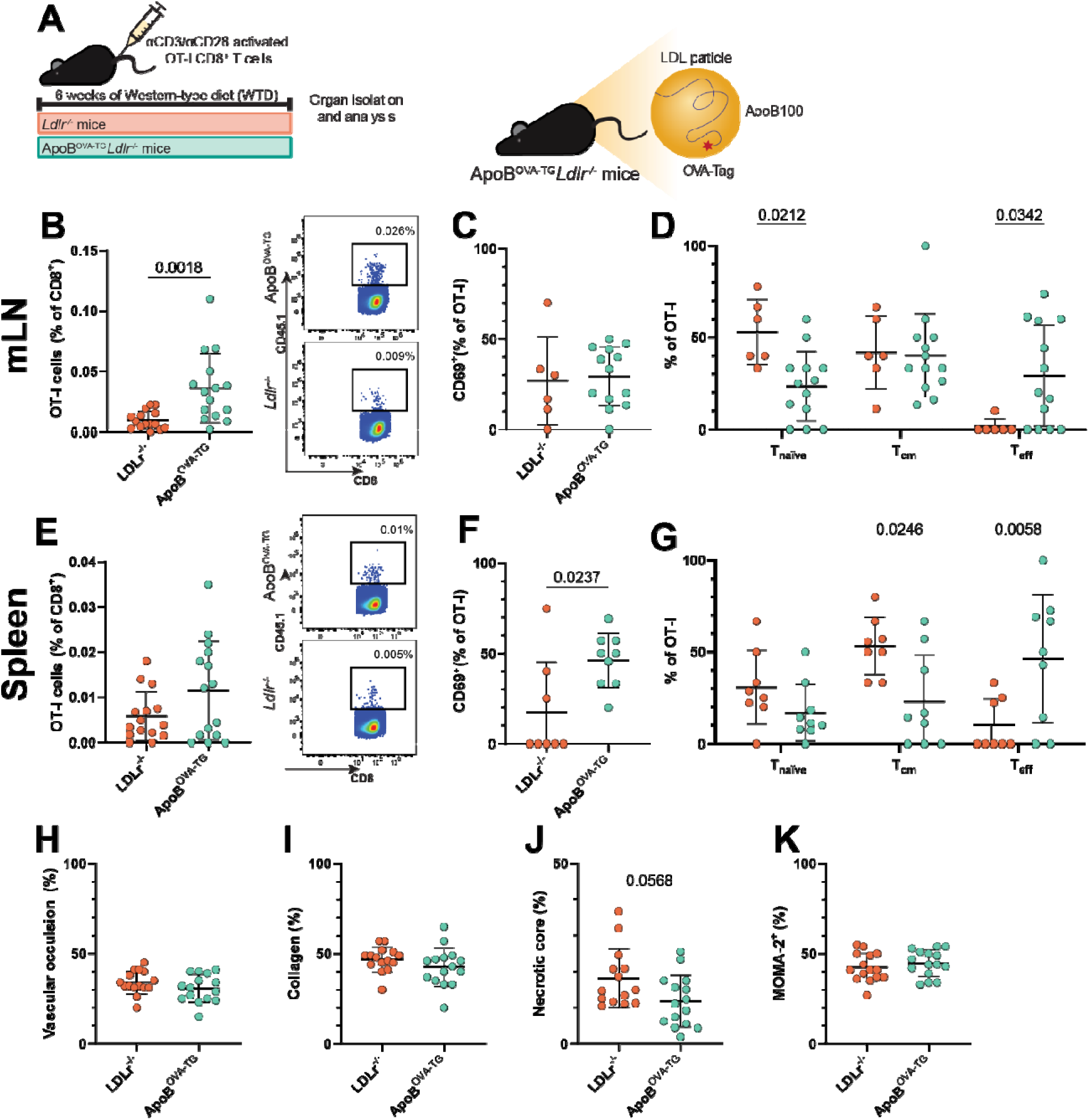
Activated OT-I circulate in antigen-specific manner towards mLN, show enhanced effector-phenotype and reduce necrotic core in Apob^OVA-TG^xLdlr^-/-^ mice. (A) Experimental setup of adoptive transfer of ex vivo activated OT-I T-cells into Ldlr^-/-^ mice or Apob^OVA-TG^xLdlr^-/^ mice. (B) Proportion of OT-I cells among CD8^+^ T-cells in the mediastinal lymph nodes (mLN), with representative flow cytometry plots. (C) Proportion of CD69^+^ cells among mLN OT-I T-cells. (D) Memory phenotype distribution mLN: naïve (T_naïve_, CD44^-^CD62L^+^), central memory (T_cm_, CD44^+^CD62L^+^) and effector memory (T_eff_, CD44^+^CD62L^-^). (E) Proportion of OT-I cells among splenic CD8^+^ T-cells, with representative flow cytometry plots. (F) Proportion of CD69^+^ cells among splenic OT-I T-cells. (G) Memory phenotype distribution of splenic OT-I cells similar to D. (H) Vascular occlusion in aortic roots, determined by Oil-Red-O staining. (I) Collagen content and (J) necrotic core size within plaques, determined by Masson’s Trichrome staining. (K) Monocyte/macrophage content within plaques, determined by MOMA-2 staining. *Outliers were removed using the ROUT test, followed by Shapiro-Wilk normality testing. Based on normality, unpaired t-tests or Mann-Whitney tests were performed for two-group comparisons, and two-way ANOVA was performed for memory phenotype analysis. n=15 mice per group. Data are shown as mean±SD*.

Despite the small number of transferred OT-I T-cells, we observed effects on atherosclerosis development that reflected the results in absence of antigen-specific T-cells (Figure 3). We observed no difference in plaque size and monocyte/macrophage content between *Apob^OVA- TG^xLdlr*^-/-^ and *Ldlr*^-/-^ mice (Figure 4H,K), but observe a clear trend towards reduced necrotic core size in the *Apob^OVA-TG^xLdlr*^-/-^ mice (Figure 4J). These results confirm a limited number of antigen-specific CD8^+^ T-cells can improve plaque stability in experimental atherosclerosis.

### ApoB-reactive CD8^+^ T-cells accumulate in plaques and reduce CD47 expression

We next sought to understand how ApoB-specific CD8^+^ T-cells can be atheroprotective. Previous work has suggested ApoB-specific CD8^+^ T-cells can interact with lipid-loaded dendritic cells.^38^ Therefore we hypothesized that OT-I T-cells may interact with lipid-loaded macrophages (foam cells) in an antigen-specific manner and eliminate them before they accumulate or become inflammatory. To understand whether OT-I T-cells accumulate in the plaque and can potentially eliminate foam cells, we adoptively transferred 6 million activated OT-I CD8^+^ T-cells into *Apob^OVA-TG^xLdlr^-/-^*mice, and compared them to control mice on the same genetic background (Figure 5A). OT-I T-cells accumulated readily within atherosclerotic plaques, with frequencies exceeded only by the liver and the circulation (Figure 5B). The particularly high hepatic accumulation is consistent with the liver being the primary site of ApoB synthesis and therefore the most likely site for MHC-I presentation of ApoB-derived epitopes. Interestingly, transfer of such a large number of antigen-specific CD8^+^ T-cells resulted in a significant reduction in plaque size, driven by decreased neutral lipid deposition (Figure 5C). Plaque size was not reduced due to perturbed lipid metabolism, as serum cholesterol levels were comparable between groups (Figure 5D) and no liver toxicity was detected (Figure 5E), confirming that the reduction in plaque size reflects a local immune-mediated effect.

**Figure 5.**
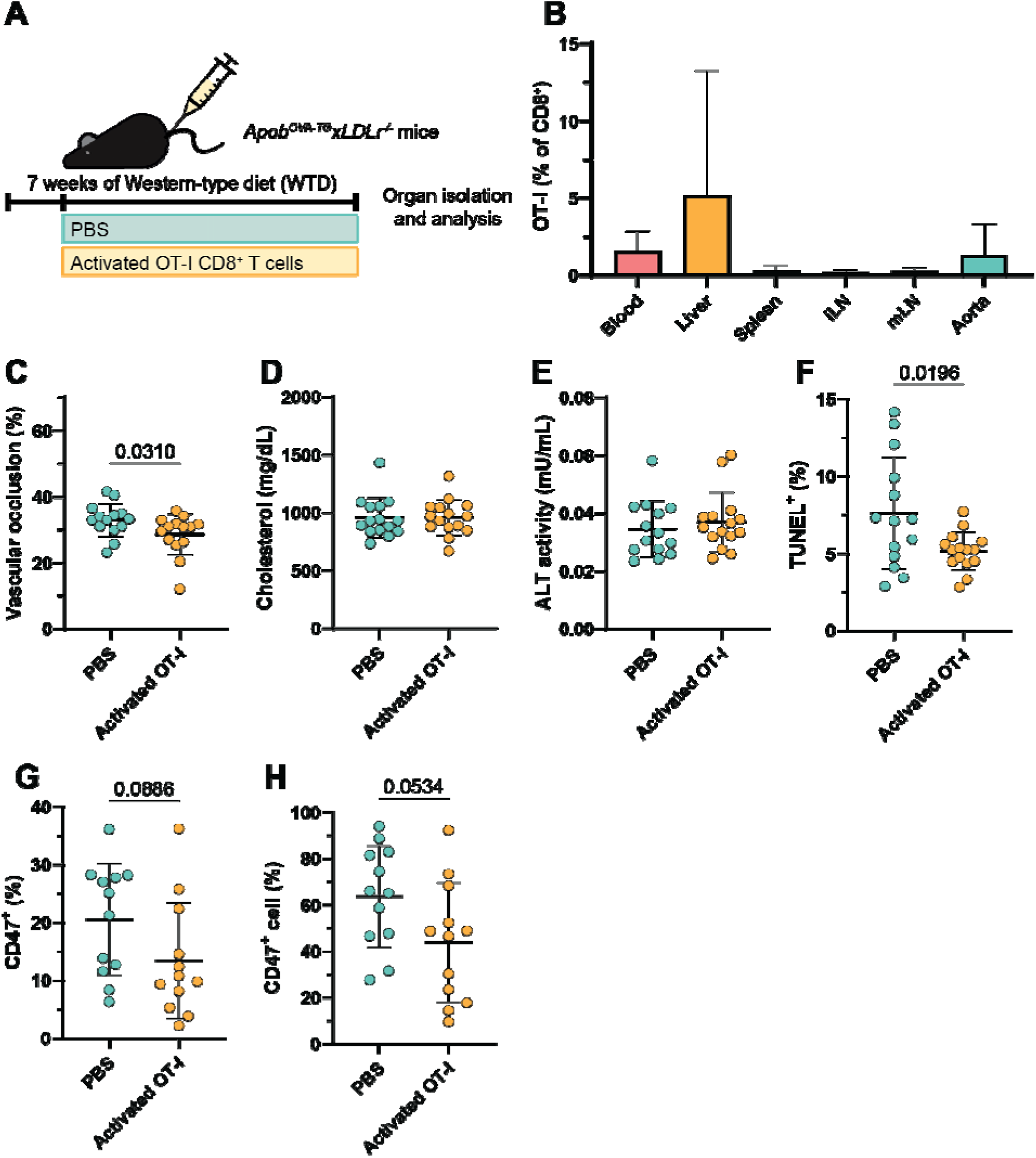
Activated atherosclerotic antigen-specific CD8^+^ T-cell transfer reduces plaque size. (A) Schematic representation of in vivo experiment where activated (6·10^6^) OT-I CD8^+^ T-cells or PBS were adoptively transferred into Apob^OVA-TG^xLdlr⁻^/^⁻ mice after one week of WTD after which the WTD was continued for another six weeks. (B) OT-I percentage of CD8^+^ T-cells per organ. (C) Vascular occlusion in aortic roots determined by Oil-Red-O staining. (D) Serum cholesterol levels. (E) Alanine Aminotransferase (ALT) activity as determined by ALT test. (F) TUNEL^+^ area in lesions as determined by TUNEL staining. (G) CD47^+^ area in lesions as determined by immunohistochemistry. (H) Percentage of CD47^+^ cells identified by cell detection. Outliers were removed using the ROUT test, followed by Shapiro-Wilk normality testing. Based *on normality, unpaired t-tests or Mann-Whitney tests were performed. n=15 mice per group. Data are shown as mean±SD*.

However, plaque composition analysis revealed no differences in collagen, necrotic core, stability index nor monocyte and macrophage content between groups (Supplementary Figure 6A-D), arguing against direct elimination of foam cells by CD8^+^ T-cells. Interestingly, TUNEL staining demonstrated significantly reduced lesional cell death in OT-I recipients (Figure 5F). As TUNEL detects fragmented genomic DNA both in apoptosis and necrosis, we next aimed to dissociate both cellular processes using the apoptosis-specific Caspase 3 staining. Remarkably, introduction of OT-I T-cells did not affect Caspase 3 levels in the plaque, and thus did not influence apoptosis (Figure S6E). Thus, antigen-specific CD8^+^ T-cells do not seem to increase apoptosis within the plaque, but do appear to affect the downstream fate of dying cells, preventing necrosis, as shown by reduced TUNEL positivity.

Apoptotic foam cells have been reported to undergo secondary necrosis when efferocytosis fails. High expression of CD47, a “don’t eat me” signal that inhibits phagocytic clearance through binding of SIRPα on neighboring phagocytes,^39^ plays a key role in preventing the removal of apoptotic foam cells from the lesion.^40^ We therefore investigated whether antigen-specific CD8^+^ T-cell interactions promote more efficient clearance of apoptotic cells by affecting CD47 expression the plaque. Immunohistochemical analysis of plaque CD47 showed a trend towards reduced CD47^+^ area in OT-I recipients (Figure 5G). Cell-based quantification of CD47 showed a similar trend upon OT-I transfer (Figure 5H). Although these observations both did not reach statistical significance, they do both point in the same direction. These findings are consistent with the possibility that antigen-specific CD8^+^ T-cells affect CD47 expression, potentially promoting the clearance of dying cells through a more efficient efferocytosis. Such a mechanism could contribute to the reduced TUNEL positivity and smaller atherosclerotic plaques observed in *Apob^OVA-TG^xLdlr^-/-^*mice receiving antigen-specific CD8^+^ T-cells.

To explore the relevance of this potential mechanism in human atherosclerosis, we next investigated CD47 expression in human atherosclerotic plaques and its association with T-cell-related gene expression. We analyzed single-cell RNA-sequencing data from carotid plaques obtained from 24 patients undergoing carotid endarterectomy surgery (GSE155512^20^, GSE159677^21^, GSE253903^22^), in which various cell types could be clustered and identified (Figure 6A, left). CITE-seq analysis revealed CD47 protein expression across all cell types (Figure, 6A right, 6B). When summing the normalized CD47 protein over all cells in each cell type, CD14^+^ myeloid cells account for the largest proportion of total CD47 expression (Figure 6C), reflecting both their CD47 expression level and number in the plaque. Thus, myeloid cells are a main source of CD47 in the atherosclerotic plaque. Given the established expression of CD47 by plaque macrophages and its potential role in cellular clearance, we next sought to investigate whether CD47 expression in human atherosclerotic lesions was associated with T-cell-related gene expression. For this, we analyzed bulk RNA-sequencing data from the Athero-Express Biobank,^29^ containing 1,093 atherosclerotic plaques (Figure 6D). *CD47* gene expression negatively correlated with *CD8A* and *CD8B* expression, suggesting that an increase in CD8 T-cell numbers reduces CD47 expression in the plaque. Importantly, positive correlations were observed for *CD47* with *CD3E* and *CD4*, showing this correlation is not merely an effect of an increase in T-cells relative to macrophages. As these associations do not establish a direct relationship between CD8^+^ T-cells and CD47 expression, we next investigated whether antigen-specific CD8^+^ T-cell interactions can directly affect CD47 expression on macrophages and foam cells *in vitro*. Bone-marrow-derived macrophages were differentiated into foam cells and co-cultured with either wildtype or OVA-responsive OT-I CD8^+^ T-cells, in the absence or presence of OVA peptide (Figure 6E). Foam cells expressed higher levels of CD47 compared to macrophages and CD47 expression was unaffected by co-culture with wildtype CD8^+^ T-cells regardless of OVA peptide loading (Figure 6F). In contrast, CD47 expression was significantly reduced on both macrophages and foam cells co-cultured with OT-I CD8^+^ T-cells, in the presence of OVA peptide. Co-culture in absence of OVA peptide had no effect on CD47 expression. These findings indicate that antigen-specific CD8^+^ T-cell interactions selectively reduce CD47 expression on antigen-presenting macrophages and foam cells, potentially promoting their efferocytotic clearance and providing a mechanistic basis for the reduced lesional cell death observed *in vivo*.

**Figure 6.**
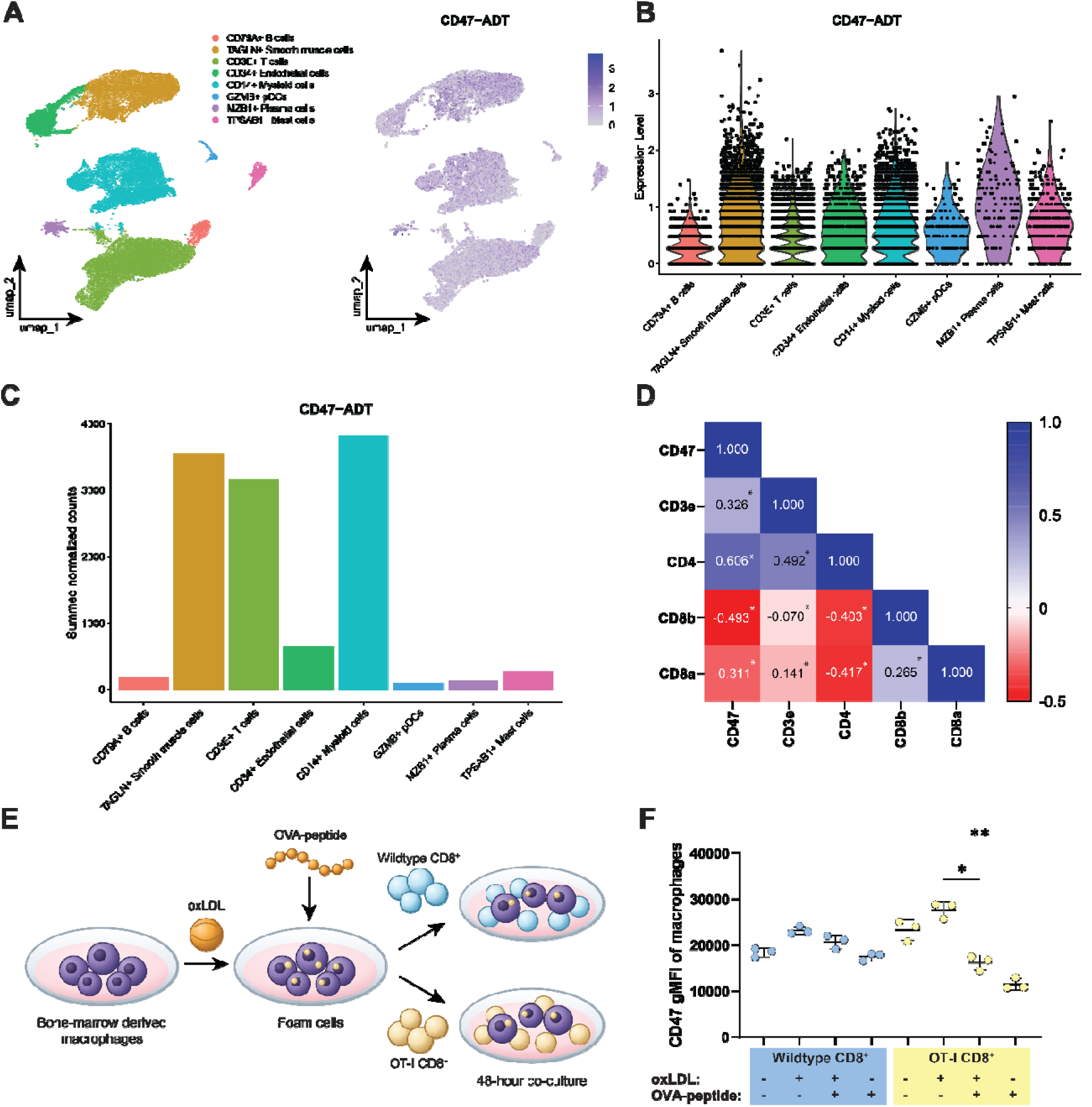
CD47 is expressed in human atherosclerotic plaques and downregulated on foam cells by antigen-specific CD8^+^ T-cells in vitro. (A) UMAP representation of cells from 24 carotid plaques, showing the annotated cell types (left) and CD47 surface protein expression measured by CITE-seq (right). (B) Violin plot of CD47 protein expression per cell type. (C) Summed normalized CD47 protein across all cells in each cell type, reflecting expression level and cell quantity. (D) Correlation heatmap determined by Spearman correlation of CD47 and T-cell genes derived from bulk RNA-seq of human atherosclerotic plaques from Athero-Express Biobank. (E) Graphical representation of bone-marrow-derived macrophage (BMDM) – CD8^+^ T-cell co-culture. BMDMs were treated with oxLDL to induce foam cell formation, after which they were loaded with ovalbumin-257 peptide antigen. Subsequently, wildtype CD8^+^ T-cells or OVA-specific CD8^+^ T-cells were co-cultured for 48 hours and marker expression was assessed by *flow cytometry. (F) Geometric mean CD47 expression in macrophages after co-culture with CD8^+^ T-cells. Kruskal-Wallis ANOVA was performed. * p<0.05, ** p<0.01*.

## Discussion

Although CD8^+^ T-cells are detected in both murine and human atherosclerotic lesions, it is unclear whether these T-cells are locally responding to antigens or are merely bystanders responding to inflammation. Here we used a reductionist approach, eliminating the capacity of all T-cells to recognize antigens, using *Ldlr*-deficient mice on a *Rag1*-deficient background, and reintroducing T-cells capable of antigen recognition (wildtype) or incapable of antigen recognition (TCR receptor transgenic). Importantly, the TCR transgenic OT-I mice donors were crossed to a *Rag1*-deficient background to eliminate the possibility of endogenous TCRs mediating antigen recognition. Our data support the notion that both plaque-specific and non-specific CD8^+^ T-cells can infiltrate atherosclerotic lesions, as both WT and OT-I CD8^+^ T-cells were found in atherosclerotic lesions. Nonetheless, antigen recognition did promote accumulation of CD8^+^ T-cells in the plaque, with accumulation of OT-I cells being reduced by 63%. This difference could not be explained by a change in size of the lesions and we observed WT CD8^+^ T-cells upregulating proteins indicative of TCR engagement, such as PD1 and CD69. The latter is a key molecule antagonizing S1PR1-driven tissue egress towards the lymph nodes,^41^ suggesting that antigen-specific interactions play an important role in retention of CD8^+^ T-cells in the plaque. Alternatively, a reduced accumulation of OT-I T-cells could be explained by reduced systemic numbers, however we observed similar frequencies of WT and OT-I T-cells in the circulation and secondary lymphoid organs. Overall, this experiment supports the hypothesis that CD8^+^ T-cells infiltrate plaques regardless of their antigen-specificity, whereas antigen recognition enhances their accumulation by increasing retention. This could explain the presence of CD8^+^ T-cells in human lesions associated with common virus infections. Cytomegaly virus specific CD8^+^ T-cells, but also Epstein-Barr or Influenza A virus (IAV) specific CD8^+^ T-cells, can reach relatively high frequencies in the circulation, thereby depositing some of the most expanded clones in the atherosclerotic plaque. Previous work suggests that IAV-specific CD8^+^ T-cells potentially recognize self-epitopes within the lesion^9^ and our data suggest that if such recognition occurs, enrichment of that particular CD8^+^ T-cell clone in the plaque relative to the circulation should be expected. Supporting this broader principle, paired single-cell TCR sequencing of blood and plaque revealed autoimmune-like features in expanded effector T-cells with preferential clonal enrichment in the lesion,^11^ consistent with antigen-driven retention shaping T-cell clonality in human atherosclerosis.

The observation that plaques contain antigen-specific and antigen-independent CD8^+^ T-cells also bring up the question what their individual contributions to disease development are. Intervening in the total CD8^+^ T-cell population in murine models has shown mixed results, with CD8^+^ T-cells reducing, contributing or having no effect on atherosclerosis progression.^12, 42^ Our results consistently show an atheroprotective effect of antigen-specific CD8^+^ T-cells. Perturbing antigen-specific CD8^+^ T-cell interactions using the *Rag1*^-/-^*Ldlr*^-/-^ model increased plaque instability, defined by less collagen deposition and increased necrotic core formation. In an opposite approach, addition of a relatively small number of ApoB-specific CD8^+^ T-cells in *Apob^OVA-TG^xLdlr*⁻*^/^*⁻ mice, decreased necrotic core size compared to introduction in *Ldlr*^-/-^ mice. These models both have limitations, but make a compelling case when combined. Whereas the *Rag1*^-/-^*Ldlr*^-/-^ model lacks B-cells, the *Apob^OVA-TG^xLdlr*⁻*^/^*⁻ does allow evaluation of antigen-specific CD8^+^ T-cell mediated effects in the presence of a functional humoral immune system. The *Apob^OVA-TG^xLdlr*⁻*^/^*⁻ model on the other hand, only allows evaluation of CD8^+^ T-cell responses to ApoB, which likely is not the only antigen CD8^+^ T-cells may respond to. The *Rag1*^-/-^*Ldlr*^-/-^ complements these results as perturbation of antigen-specific CD8^+^ T-cell response is independent of the antigen. Furthermore, the *Rag1*^-/-^*Ldlr*^-/-^ model does not allow tracking of antigen-specific CD8^+^ T-cells or interrogation of their function, limitations that can be directly addressed with the *Apob^OVA-TG^xLdlr^-/-^* model. By transferring a large number OT-I T-cells into these mice we are able to track their accumulation and activation to the atherosclerotic plaque and observed a significant reduction in neutral lipid accumulation in the plaque. *In vitro* co-culture experiments demonstrate that antigen-specific interactions between OT-I CD8^+^ T-cells and foam cells reduce CD47 expression on macrophages, suggesting that ApoB-specific CD8^+^ T-cells may prime ApoB-presenting macrophages for phagocytic clearance within the lesion, potentially reducing plaque progression. However this cytotoxic capacity has not yet been reflected *in vivo*, as MOMA-2 staining revealed no difference in macrophage content between groups. This discrepancy likely reflects the complex plaque microenvironment compared to the controlled co-culture system. Within the lesion, antigen-specific CD8^+^ T-cells represent only a small fraction of infiltrating cells and continuous monocyte recruitment replenishes the macrophage pool, potentially masking the local cytotoxic effects on overall macrophage numbers. Furthermore, CD47-mediated regulation of phagocytosis may influence the clearance of individual cells without necessarily affecting overall macrophage abundance. Thus, whether CD47 downregulation observed *in vitro* translates into enhanced efferocytosis and reduced macrophage accumulation *in vivo* therefore warrants further investigation. Human atherosclerotic lesions however, do point to CD8^+^ T-cell accumulation reducing CD47 expression in the plaque.

It should be noted that the mechanism described above is dependent on ApoB antigen presentation and may not apply to other MHC-I-restricted antigens driving CD8^+^ T-cell responses in the plaque. Indeed, we have previously shown that ApoB-derived epitopes are only a minor fraction of the HLA-I restricted antigen presented in human atherosclerotic plaques,^17^ suggesting other antigens may be engaged by CD8^+^ T-cells in the plaque. Another complicating factor in this model is the accumulation of ApoB-specific CD8^+^ T-cells in the liver. As the primary site of VLDL/LDL assembly, hepatocytes synthesize ApoB_100_,^43^ making it a major potential source of MHC-I-mediated antigen presentation. This raises the possibility that at least part of the antigen-specific CD8^+^ T-cell activation observed reflects unwanted hepatic responses. Interestingly, we did not observe signs of liver distress, judged by liver enzyme and cholesterol levels. Hepatic priming has been strongly linked to the induction of immune tolerance or early exhaustion instead of full effector differentiation, due to the liver’s role in maintaining peripheral tolerance to circulating antigens.^44^ Nonetheless, the potential interaction of ApoB-specific CD8^+^ T-cells would be a major impediment to therapeutic exploitation of this antigen.

Future work integrating single-cell transcriptomic profiling of plaque-derived and circulating CD8^+^ T-cells, combined with functional validation will be important to fully understand the mechanism by which antigen-specific CD8^+^ T-cells influence plaque progression. Our findings support a model in which antigen recognition drives CD8^+^ T-cell retention and effector differentiation within the lesion. Collectively, these data affirm antigen-specific CD8^+^ T-cells as active modulators of atherosclerotic disease and highlight their potential as a novel target for immune-based therapeutic strategies aimed at promoting plaque stability and limiting cardiovascular risk.

## Data Availability

The data supporting the findings of this study are available from the corresponding author upon reasonable request.

## Acknowledgements

We would like to thank P.J. van Santbrink for performing all genotyping analyses.

## References

1. Lindstrom M, DeCleene N, Dorsey H, Fuster V, Johnson CO, LeGrand KE, et al. Global Burden of Cardiovascular Diseases and Risks Collaboration, 1990-2021. J Am Coll Cardiol 2022;80(25):2372–2425.

2. Riaz H, Khan SU, Lateef N, Talluri S, Khan MS, Desai MY. Residual inflammatory risk after contemporary lipid lowering therapy. Eur Heart J Qual Care Clin Outcomes 2020;6(2):105–111.

3. Ridker Paul M, Everett Brendan M, Thuren T, MacFadyen Jean G, Chang William H, Ballantyne C, et al. Antiinflammatory Therapy with Canakinumab for Atherosclerotic Disease. New England Journal of Medicine;377(12):1119–1131.

4. Ridker PM, MacFadyen JG, Thuren T, Libby P. Residual inflammatory risk associated with interleukin-18 and interleukin-6 after successful interleukin-1β inhibition with canakinumab: further rationale for the development of targeted anti-cytokine therapies for the treatment of atherothrombosis. Eur Heart J 2020;41(23):2153–2163.

5. Hinkley H, Counts DA, VonCanon E, Lacy M. T Cells in Atherosclerosis: Key Players in the Pathogenesis of Vascular Disease. Cells 2023;12(17).

6. Saigusa R, Winkels H, Ley K. T cell subsets and functions in atherosclerosis. Nature Reviews Cardiology 2020;17(7):387–401.

7. Schäfer S, Zernecke A. CD8(+) T Cells in Atherosclerosis. Cells 2020;10(1).

8. Smit V, de Mol J, Schaftenaar FH, Depuydt MAC, Postel RJ, Smeets D, et al. Single-cell profiling reveals age-associated immunity in atherosclerosis. Cardiovasc Res 2023;119(15):2508–2521.

9. Chowdhury RR, D’Addabbo J, Huang X, Veizades S, Sasagawa K, Louis DM, et al. Human Coronary Plaque T Cells Are Clonal and Cross-React to Virus and Self. Circ Res 2022;130(10):1510–1530.

10. Depuydt MAC, Prange KHM, Slenders L, Örd T, Elbersen D, Boltjes A, et al. Microanatomy of the Human Atherosclerotic Plaque by Single-Cell Transcriptomics. Circ Res 2020;127(11):1437–1455.

11. Depuydt MAC, Schaftenaar FH, Prange KHM, Boltjes A, Hemme E, Delfos L, et al. Single-cell T cell receptor sequencing of paired human atherosclerotic plaques and blood reveals autoimmune-like features of expanded effector T cells. Nat Cardiovasc Res 2023;2(2):112–125.

12. Kyaw T, Winship A, Tay C, Kanellakis P, Hosseini H, Cao A, et al. Cytotoxic and proinflammatory CD8+ T lymphocytes promote development of vulnerable atherosclerotic plaques in apoE-deficient mice. Circulation 2013;127(9):1028–39.

13. Jorgacevic I, Shaikh H, Ali H, Bundalo M, Schäfer S, Kern MAG, et al. Allogeneic haematopoietic cell transplantation promotes atherosclerosis in mice via CD8+ T cells. Cardiovasc Res 2025;121(17):2668–2678.

14. van Duijn J, de Jong MJM, Benne N, Leboux RJT, van Ooijen ME, Kruit N, et al. Tc17 CD8+ T cells accumulate in murine atherosclerotic lesions, but do not contribute to early atherosclerosis development. Cardiovasc Res 2021;117(14):2755–2766.

15. Wang Y, Li P, Li W, Zou Y, Zhao Y, Ma L, et al. CD80(+) Macrophage-Induced IL-17(+)CD8(+) T cells accumulate in hyperlipidemic patients and murine vascular lesions to promote atherosclerotic progression. Cell Commun Signal 2026;24(1).

16. Stelma F, de Niet A, Sinnige MJ, van Dort KA, van Gisbergen KPJM, Verheij J, et al. Human intrahepatic CD69 + CD8+ T cells have a tissue resident memory T cell phenotype with reduced cytolytic capacity. Scientific Reports 2017;7(1):6172.

17. de Jong MJM, Schaftenaar FH, Depuydt MAC, Lozano Vigario F, Janssen GMC, Peeters J, et al. Virus-Associated CD8(+) T-Cells Are Not Activated Through Antigen-Mediated Interaction Inside Atherosclerotic Lesions. Arterioscler Thromb Vasc Biol 2024;44(6):1302–1314.

18. Wang H, Yang H, Shivalila CS, Dawlaty MM, Cheng AW, Zhang F, et al. One-step generation of mice carrying mutations in multiple genes by CRISPR/Cas-mediated genome engineering. Cell 2013;153(4):910–8.

19. Bankhead P, Loughrey MB, Fernández JA, Dombrowski Y, McArt DG, Dunne PD, et al. QuPath: Open source software for digital pathology image analysis. Scientific Reports 2017;7(1):16878.

20. Pan H, Xue C, Auerbach BJ, Fan J, Bashore AC, Cui J, et al. Single-Cell Genomics Reveals a Novel Cell State During Smooth Muscle Cell Phenotypic Switching and Potential Therapeutic Targets for Atherosclerosis in Mouse and Human. Circulation 2020;142(21):2060–2075.

21. Alsaigh T, Evans D, Frankel D, Torkamani A. Decoding the transcriptome of calcified atherosclerotic plaque at single-cell resolution. Commun Biol 2022;5(1):1084.

22. Bashore AC, Yan H, Xue C, Zhu LY, Kim E, Mawson T, et al. High-Dimensional Single-Cell Multimodal Landscape of Human Carotid Atherosclerosis. Arterioscler Thromb Vasc Biol 2024;44(4):930–945.

23. Hao Y, Stuart T, Kowalski MH, Choudhary S, Hoffman P, Hartman A, et al. Dictionary learning for integrative, multimodal and scalable single-cell analysis. Nature Biotechnology 2024;42(2):293–304.

24. Germain PL, Lun A, Garcia Meixide C, Macnair W, Robinson MD. Doublet identification in single-cell sequencing data using scDblFinder. F1000Res 2021;10:979.

25. Yin Y, Yajima M, Campbell JD. Characterization and decontamination of background noise in droplet-based single-cell protein expression data with DecontPro. Nucleic Acids Research 2024;52(1):e4–e4.

26. Kim H, Park I, Park J-E, Kim JK, Seo M, Kim JK. scICE: enhancing clustering reliability and efficiency of scRNA-seq data with multi-cluster label consistency evaluation. Nature Communications 2025;16(1):6031.

27. Kim H, Chang W, Chae SJ, Park J-E, Seo M, Kim JK. scLENS: data-driven signal detection for unbiased scRNA-seq data analysis. Nature Communications 2024;15(1):3575.

28. Wickham H. ggplot2: Elegant Graphics for Data Analysis: Springer-Verlag New York; 2016.

29. Verhoeven BAN, Velema E, Schoneveld AH, Vries JPPMd, de Bruin P, Seldenrijk CA, et al. Athero-express: Differential atherosclerotic plaque expression of mRNA and protein in relation to cardiovascular events and patient characteristics. Rationale and design. European Journal of Epidemiology 2004;19(12):1127–1133.

30. Parma L, Sachs N, Sobczak N, Li Z, Merchant K, Pauli J, et al. CXCL12 Derived From ACKR1+ Intraplaque Neovessels Mediates CD8+ T Cell Recruitment in Human Atherosclerosis. Circulation 2025;151(8):581–584.

31. Mulholland M, Chalou A, Andersson SHA, Depuydt MAC, Yu Y, Lin S, et al. Progenitor exhausted PD-1+ T cells are cellular targets of immune checkpoint inhibition in atherosclerosis. Nature Cardiovascular Research 2025;4(10):1311–1328.

32. Sad S, Marcotte R, Mosmann TR. Cytokine-induced differentiation of precursor mouse CD8+ T cells into cytotoxic CD8+ T cells secreting Th1 or Th2 cytokines. Immunity 1995;2(3):271–9.

33. Schüler T, Kammertoens T, Preiss S, Debs P, Noben-Trauth N, Blankenstein T. Generation of tumor-associated cytotoxic T lymphocytes requires interleukin 4 from CD8(+) T cells. J Exp Med 2001;194(12):1767–75.

34. von Vietinghoff S, Ley K. Interleukin 17 in vascular inflammation. Cytokine Growth Factor Rev 2010;21(6):463–9.

35. Lozano Vigario F, Molenaar J, Simó Vesperinas I, van der Zon M, Crone NSA, de Jong MJM, et al. Immunopeptidomics analysis of human atherosclerosis plaques identifies antigenic drivers of atherosclerosis. Atherosclerosis 2025;409:120509.

36. Hermansson A, Ketelhuth DF, Strodthoff D, Wurm M, Hansson EM, Nicoletti A, et al. Inhibition of T cell response to native low-density lipoprotein reduces atherosclerosis. J Exp Med 2010;207(5):1081–93.

37. Roy P, Parashar S, Sidney J, Mehta M, Tandon I, Beddall M, et al. Increased Autoreactive CD8(+) T Cells to Apolipoprotein B Epitopes in Patients With Severe Coronary Artery Disease. Circ Res 2026;139(6):e326804.

38. Chyu KY, Zhao X, Dimayuga PC, Zhou J, Li X, Yano J, et al. CD8+ T cells mediate the athero-protective effect of immunization with an ApoB-100 peptide. PLoS One 2012;7(2):e30780.

39. Bess SN, Igoe MJ, Muldoon TJ. The Physiological and Therapeutic Role of CD47 in Macrophage Function and Cancer. Immunol Invest 2025;54(1):112–146.

40. Kojima Y, Volkmer JP, McKenna K, Civelek M, Lusis AJ, Miller CL, et al. CD47-blocking antibodies restore phagocytosis and prevent atherosclerosis. Nature 2016;536(7614):86–90.

41. Walsh DA, Borges da Silva H, Beura LK, Peng C, Hamilton SE, Masopust D, et al. The Functional Requirement for CD69 in Establishment of Resident Memory CD8+ T Cells Varies with Tissue Location. The Journal of Immunology 2019;203(4):946–955.

42. Cochain C, Koch M, Chaudhari SM, Busch M, Pelisek J, Boon L, et al. CD8+ T Cells Regulate Monopoiesis and Circulating Ly6C-high Monocyte Levels in Atherosclerosis in Mice. Circ Res 2015;117(3):244–53.

43. Edge SB, Hoeg JM, Schneider PD, Brewer HB, Jr. Apolipoprotein B synthesis in humans: Liver synthesizes only apolipoprotein B-100. Metabolism - Clinical and Experimental 1985;34(8):726–730.

44. Bowen DG, Zen M, Holz L, Davis T, McCaughan GW, Bertolino P. The site of primary T cell activation is a determinant of the balance between intrahepatic tolerance and immunity. J Clin Invest 2004;114(5):701–12.

