## Supplemental Figures and Tables for "CD8^+^ T-cells protect against experimental atherosclerosis through antigen-specific interactions in the plaque"

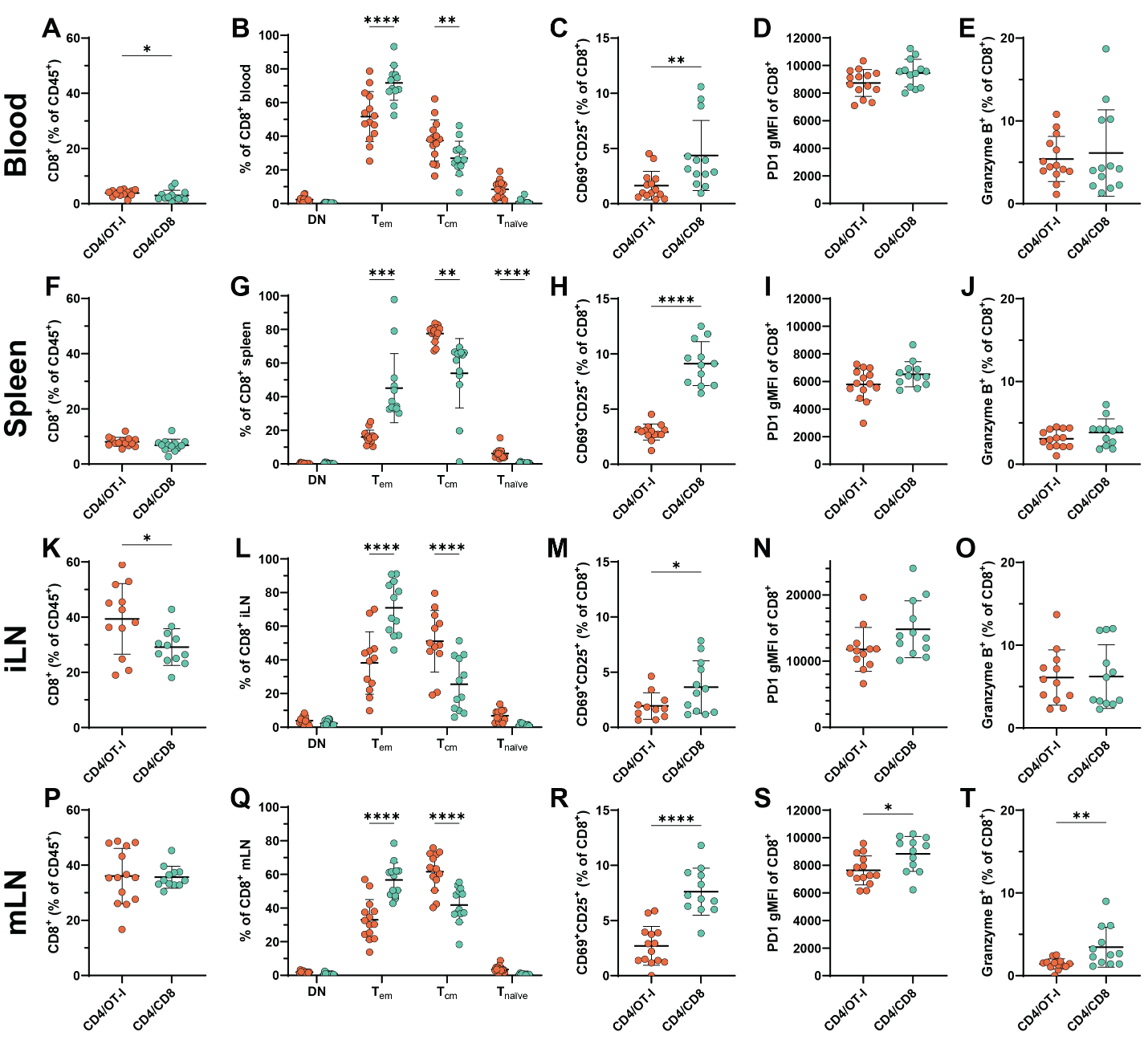

***Supplemental Figure 1. Transferred CD8^+^ T cells originating from wildtype background show more effector-like activation compared to antigen-restricted CD8^+^ T cells systemically.*** *The proportion of (A) CD8^+^ T cells of immune cells, (B) CD8^+^ T-cell phenotype with double negative (DN, CD44^-^CD62L^-^), effector-memory (T_em_, CD44^+^CD62L^-^), central-memory (T_cm_, CD44^+^CD62L^+^) and naïve (T_naïve_, CD44^-^CD62L^+^), (C) CD25^+^CD69^+^ CD8^+^ T cells, (D) PD1 gMFI of CD8^+^ T cells, and (E) Granzyme B^+^ of CD8^+^ T cells in the blood (A-E), spleen (F-J), inguinal lymph node (iLN) (K-O) and mediastinal lymph node (mLN) (P-T) after adoptive transfer into Rag^-/-^Ldlr^-/-^ mice on 10 weeks of Western-type diet. Outliers were removed using the ROUT test, followed by Shapiro-Wilk normality testing. Based on normality, unpaired t-tests or Mann-Whitney tests were performed. n=13-14 mice per group. Data are shown as mean ± SD*.*p<0.05, **p<0.01, ***p<0.001, ****p<0.0001.

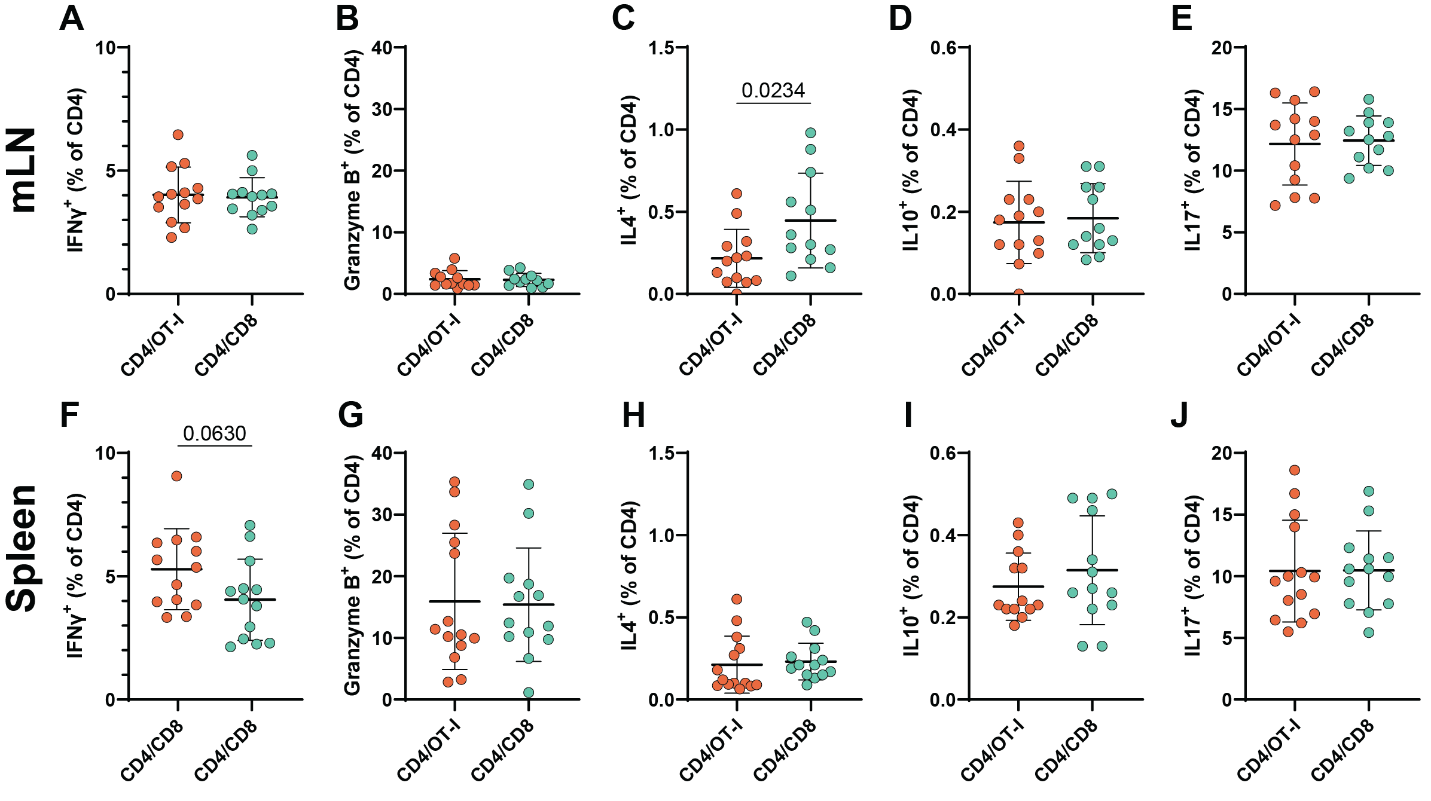

**Supplemental Figure 2.** ***Antigen-specific CD8^+^ T-cell transfer does not affect CD4 T-cell cytokine production****. Percentage of CD4^+^ T cells producing cytokines and granzymes in mediastinal lymph nodes (mLN) (A-E) and spleen (F-J) after 6 hours of ex vivo PMA/ionomycin stimulation. Interferon gamma (A,F), Granzyme B (B,G), interleukin-4 (C,H), interleukin-10 (D,I) and interleukin-17 production (E,J) in CD8^+^ T cells. Outliers were removed using the ROUT test, followed by Shapiro-Wilk normality testing. Based on normality, unpaired t-tests or Mann-Whitney tests were performed. n=13-14 mice per group. Data are shown as mean ± SD.*

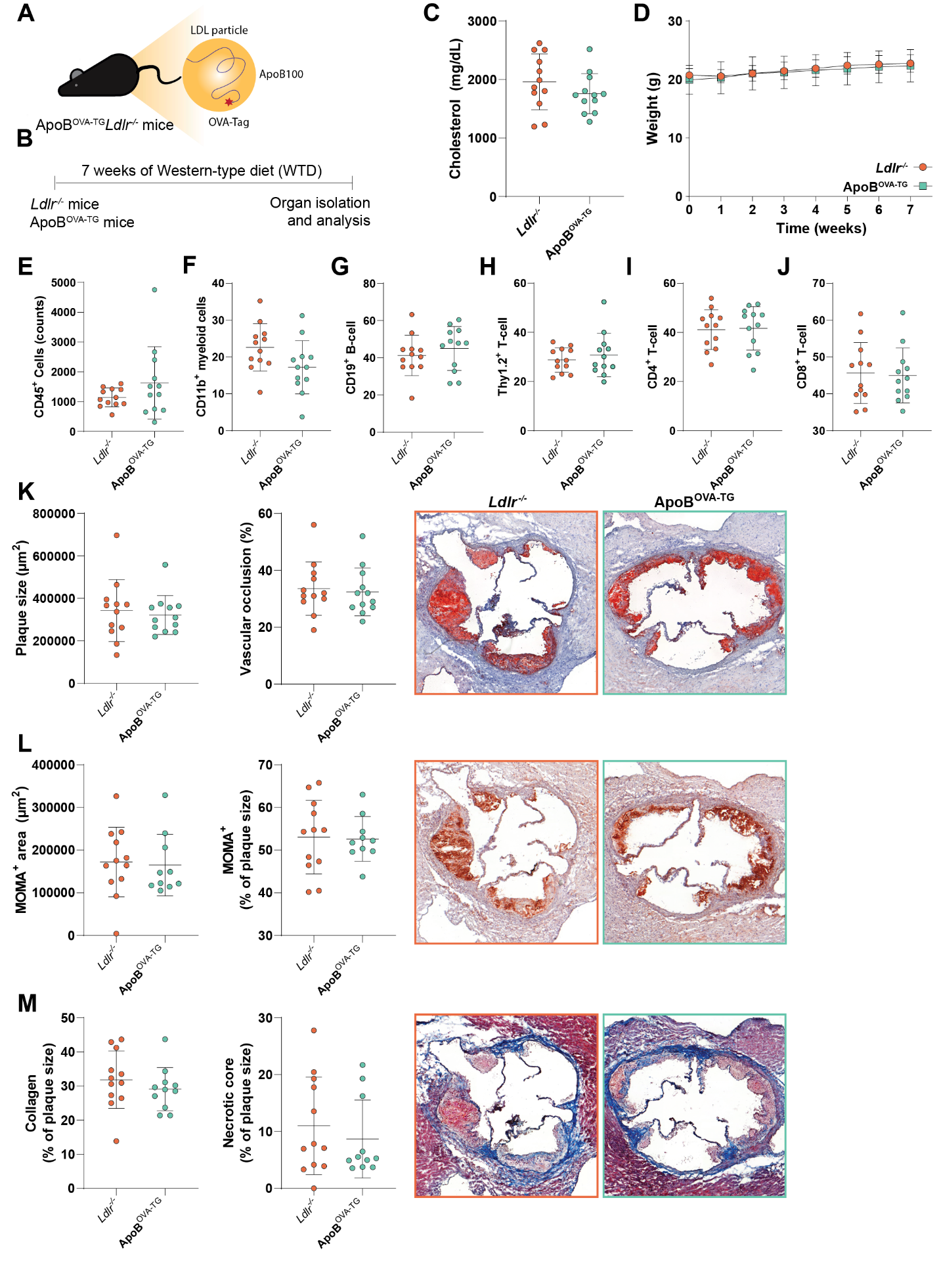

***Supplemental Figure 3.*** ***Vitality and atherosclerotic lesion development of Apob^OVA-TG^x Ldlr^-/-^ mice, in comparison to Ldlr^-/-^ mice.*** *(A) Graphical representation of Apob^OVA-TG^xLdlr^-/-^ mice. (B) Experimental design for assessing plaque progression in Apob^OVA-TG^xLdlr^-/-^ mice. (C) Plasma cholesterol levels of the mice during western type diet. (D) Mouse weight of Apob^OVA-TG^xLdlr^-/-^ and Ldlr^-/-^ mice in weeks, during 7 weeks of Western type diet. The number of (E) CD45^+^ leukocytes and the proportion of (F) CD11b^+^ myeloid cells, (G) CD19^+^ B cells, and (H) Thy1.2^+^ T cells of CD45^+^ cells, and (I) CD4^+^ T cells, and (J) CD8^+^ T cells of Thy1.2^+^ T cells in the aorta after 7 weeks of western type diet. (K) Quantification of plaque size and vascular stenosis in the trivalve area of the heart, with representative images from Ldlr^-/-^ and Apob^OVA-TG^xLdlr^-/-^ mice. (L) Evaluation of absolute and relative monocyte/macrophage positive area in the lesion, with representative images. (M) Assessment of collagen content and necrotic core size, determined by Masson’s Trichrome staining, including corresponding images. Outliers were removed using the ROUT test, followed by Shapiro-Wilk normality testing. Based on normality, unpaired t-tests or Mann-Whitney tests were performed. n=13-14 mice per group. Data are shown as mean ± SD.*

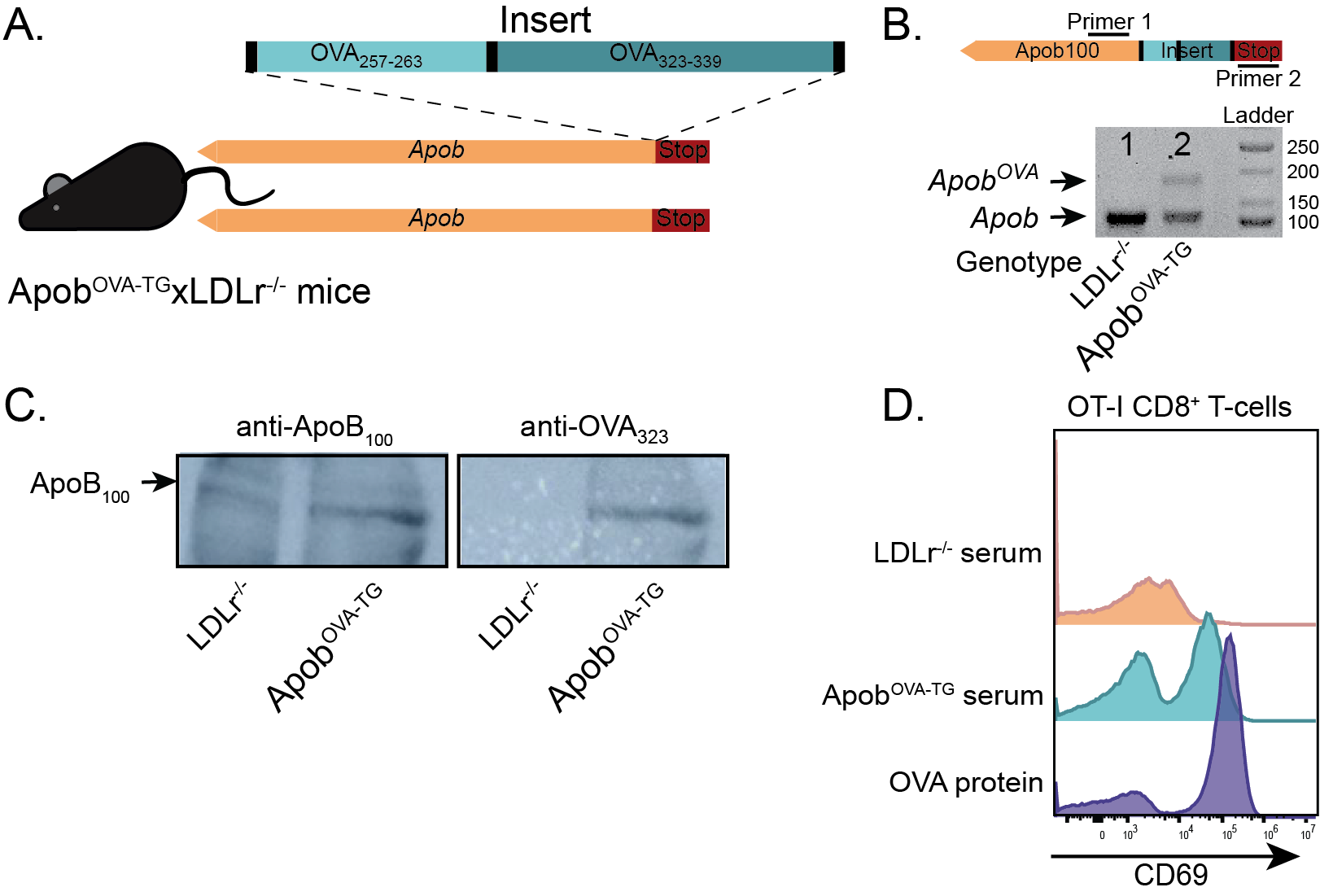

***Supplemental Figure 4. Generation and validation of Apob^OVA-TG^xLdlr^-/-^ mice, and serum-induced activation of OT-I CD8^+^ T cells***. *(A) Schematic of the OVA gene cassette inserted at the C-terminus of the Apob gene. (B) Genotyping of Apob^OVA-TG^xLdlr^-/-^ mice. (C) Western blot confirming fusion of OVA_323_ to ApoB, detected with anti-ApoB_100_ and anti-OVA_323_ antibodies. (D) OT-I splenocytes were pulsed overnight with serum derived from Ldlr^-/-^ or Apob^OVA-TG^xLdlr^-/-^ mice. CD69 expression on OT-I CD8^+^ T cells was assessed by flow cytometry after 24h.*

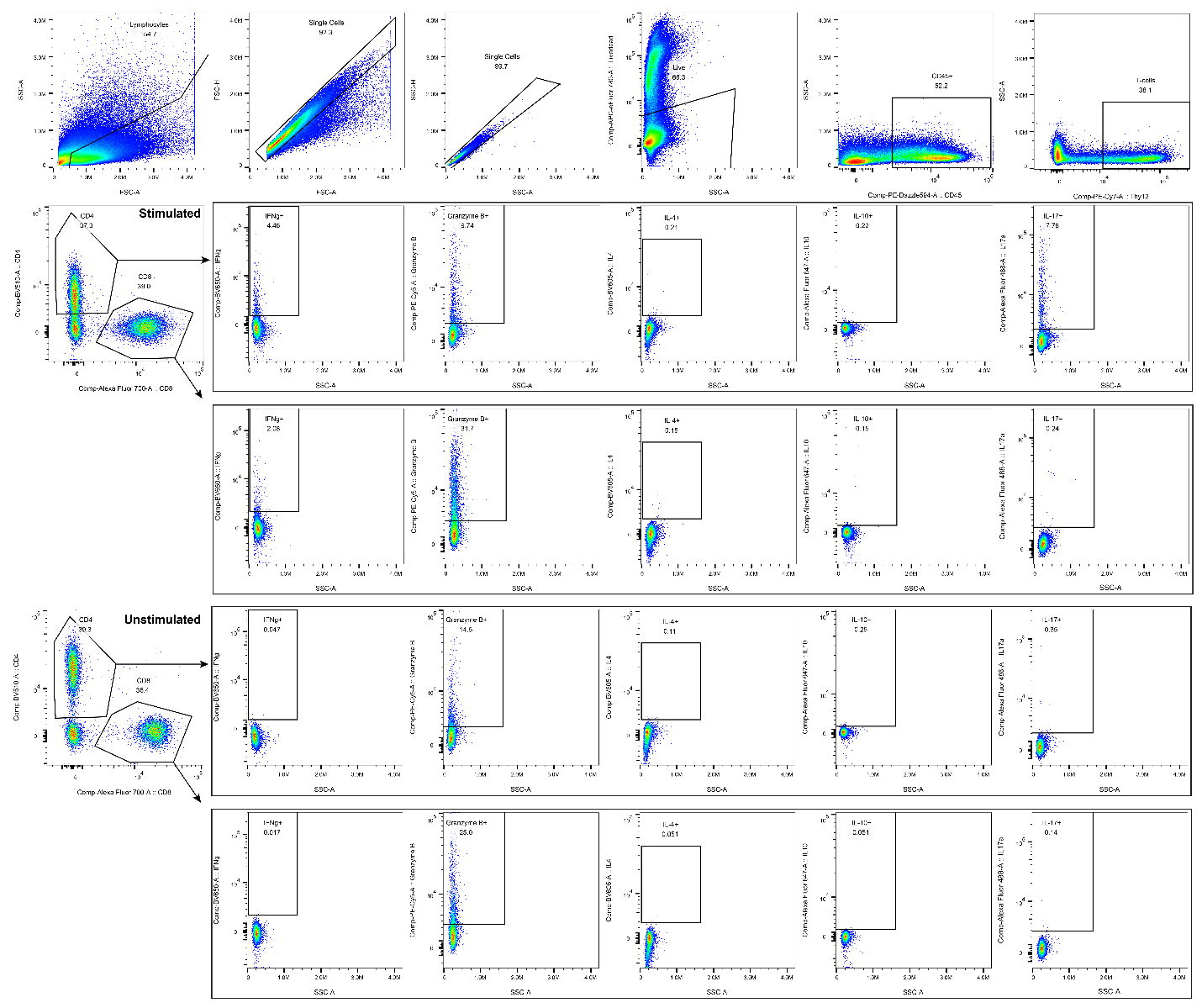

***Supplemental Figure 5. Flow cytometry gating strategy for the identification of cytokine-expressing CD4^+^ and CD8^+^ T cells in Rag1^-/-^Ldlr^-/-^ following adoptive transfer of WT or OT-I T cells and ex vivo stimulation with PMA/ionomycin.*** *Live CD45^+^Thy1.2^+^ T cells were gated into CD4^+^ and CD8^+^ subsets, followed by assessment of cytokine expression. Representative plots are shown from one mouse with a sufficient T-cell population for analysis, together with an unstimulated control mouse sample.*

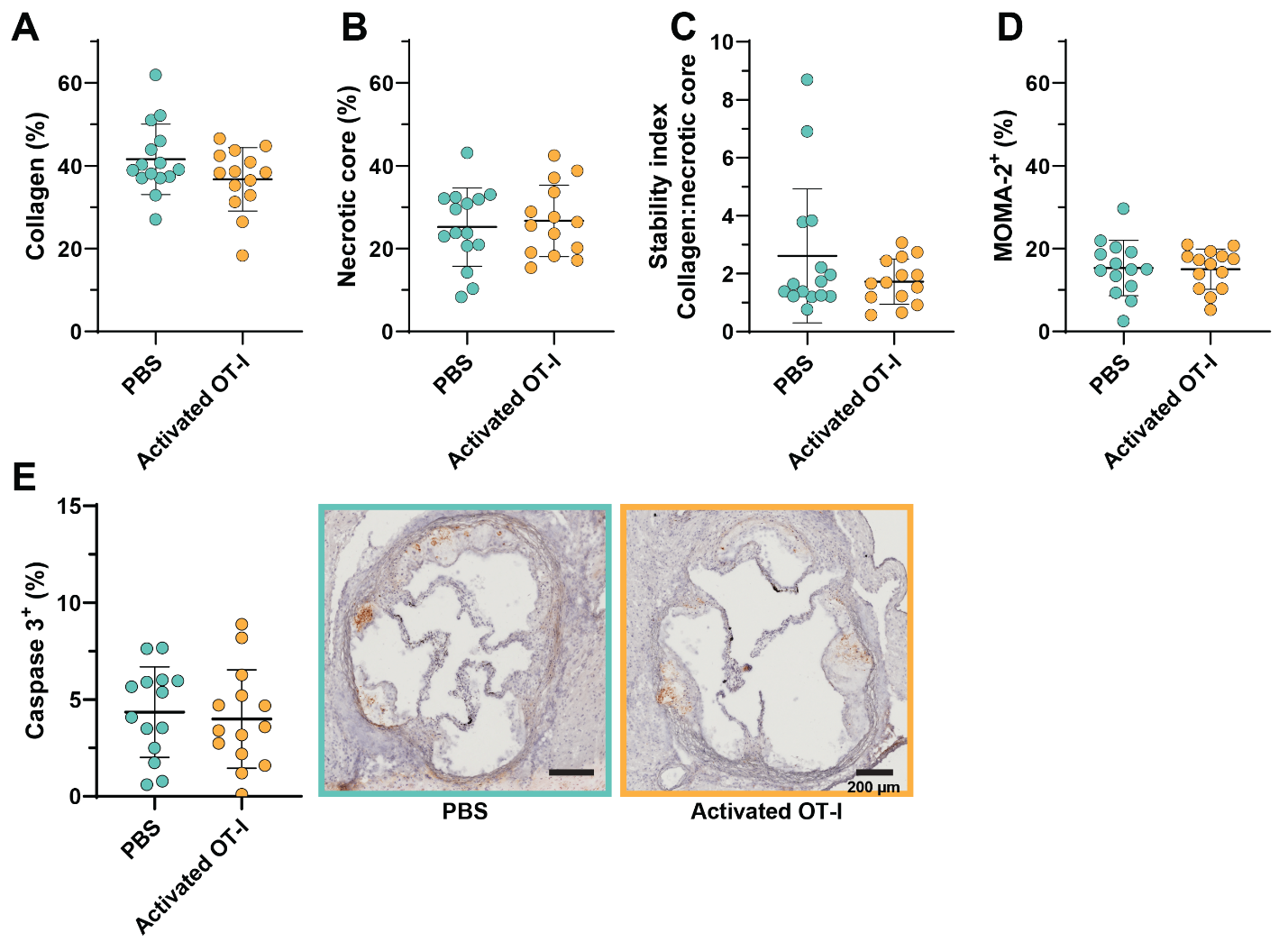
***Supplemental Figure 6. Activated atherosclerotic antigen-specific CD8^+^ T-cell transfer has no effect on plaque stability, monocyte/macrophage content or Caspase 3 expression in the lesion.*** *Activated (6∙10^6^)* *OT-I CD8^+^ T cells* *or PBS were adoptively transferred into Apob^OVA-TG^xLdlr⁻^/^⁻ mice after one week of WTD after which the WTD was continued for another six weeks. (A) Collagen content and (B) necrotic core size within plaques, determined by Masson’s Trichrome staining. (C) Stability index of plaques as determined by ratio of collagen per necrosis. (D) Monocyte/macrophage content within plaques, determined by MOMA-2 staining. (E) Caspase-3^+^ area in aortic root plaques as determined by immunohistochemistry, with representative images.*

### Supplementary tables

***Supplementary Table 1.*** *Antibody list for flow cytometry.*

| **Marker** | **Fluorochrome** | **Clone** | **Supplier** | **Catalog number** | **Dilution** |
| --- | --- | --- | --- | --- | --- |
| CD11b | PE-Cy5 | M1/70 | BioLegend | 101209 | 1000 |
| CD11b | PerCP-Cy5.5 | M1/70 | BioLegend | 101227 | 1000 |
| CD11c | PerCP-Cy5.5 | N418 | BioLegend | 117328 | 500 |
| CD19 | PE-Cy7 | 6D5 | BioLegend | 115519 | 1000 |
| CD25 | PE | PC61.5 | eBioscience | 12-0251-83 | 2000 |
| CD25 | Alexa Fluor 488 | PC61.5 | BioLegend | 102017 | 2000 |
| CD4 | BV510 | RM4-5 | BioLegend | 100559 | 1000 |
| CD44 | FITC | IM7 | eBioscience | 11-0441-85 | 500 |
| CD44 | PE-Dazzle 594 | IM7 | BioLegend | 103056 | 1000 |
| CD44 | PerCP | IM7 | BioLegend | 103035 | 500 |
| CD45 | PE-Dazzle 594 | 30-F11 | BioLegend | 103146 | 4000 |
| CD45 | APC-Fire 750 | 30-F11 | BioLegend | 103154 | 1000 |
| CD45.1 | eFluor450 | A20 | eBioscience | 48-0453-80 | 500 |
| CD47 | PE-Cy7 | miap301 | BioLegend | 127523 | 1000 |
| CD62L | BV570 | MEL-14 | BioLegend | 104433 | 500 |
| CD62L | BV605 | MEL-14 | BioLegend | 104438 | 500 |
| CD69 | BV785 | H1.2F3 | BioLegend | 104543 | 500 |
| CD69 | FITC | H1.2F3 | BioLegend | 104532 | 500 |
| CD8a | Alexa Fluor 700 | 53-6.7 | BioLegend | 100730 | 1000 |
| CD8a | APC-Fire 810 | 53-6.7 | BioLegend | 104008 | 1000 |
| CD8a | PE-Dazzle 594 | 53-6.7 | BioLegend | 100761 | 500 |
| F4/80 | BV711 | BM8 | BioLegend | 123147 | 500 |
| Granzyme B | PE-Cy5 | QA16A02 | BioLegend | B423437 | 250 |
| IFNγ | BV650 | XMG1.2 | BioLegend | 505832 | 500 |
| IL10 | AF647 | JES5-16E3 | BioLegend | 505014 | 500 |
| IL17A | Alexa Fluor 488 | TC11-18H10.1 | BioLegend | 506909 | 500 |
| IL4 | BV605 | 11B11 | BioLegend | 504125 | 500 |
| PD-1 | PE-Fire 810 | 29F.1A12 | BioLegend | 135253 | 1000 |
| Thy1.2 | PE-Cy7 | 53-2.1 | BioLegend | 140310 | 1500 |
| TruStain FcX™ |  | 93 | BioLegend | 101320 | 250 |
| Fixable Viability Dye | eFluor 780 |  | eBioscience | 65-0865-18 | 4000 |
| Zombie NIR Fixable Viability Kit | Zombie NIR |  | BioLegend | 423106 | 4000 |
